# Effective prediction of IL-17 inducing peptides using hybrid approach: iIL17pred

**DOI:** 10.1101/2025.01.08.631619

**Authors:** Pooja Arora, Neha Periwal, Lakshay Aggarwal, Vikas Sood, Baljeet Kaur

## Abstract

**Background:** Interleukin 17 (IL-17) plays a crucial role in regulating the immune system and is associated with numerous diseases. Modulating IL-17 levels has demonstrated potential in mitigating disease symptoms, positioning it as a compelling target for drug development. Therefore, identifying and characterizing novel drug molecules capable of influencing IL-17 levels is critical. Recent advances in therapeutic peptides underscore their promise as attractive drug candidates, inspiring the development of IL-17 modulating peptides.

**Results:** This study aimed to enhance existing methods for efficiently classifying IL-17 inducing peptides. Positive and negative datasets were obtained from the Immune Epitope Database, and peptide features were extracted using the pfeature algorithm. A three-stage hybrid approach combining BLAST, MERCI, and machine learning techniques was employed to accurately classify IL-17-inducing peptides.

**Conclusions:** Extensive benchmarking experiments revealed that our proposed method outperforms existing techniques i.e IL17eScan, across key performance metrics, including sensitivity, accuracy, and area under the curve-receiver operating characteristics (AUC-ROC). Our algorithm achieved an accuracy of 88.08% and a Matthews Correlation Coefficient (MCC) of 0.68 on an external dataset, significantly surpassing the accuracy of 78.57% and MCC of 0.57 achieved by existing methods under comparable conditions. This demonstrates a substantial improvement in IL-17 peptide classification. The results are accessible via a user-friendly web server (http://www.soodlab.com/iil17pred/). These findings hold significant potential for predicting IL-17-inducing peptides, which can be further validated experimentally.

## Introduction

Interleukins are a type of cytokines that were initially thought to be produced from the leukocytes. However, further studies proved that these molecules are secreted from a vast variety of cells. Primarily these molecules are known for the activation and differentiation of immune cells thereby making them critical players in the body’s immune responses. Among the various interleukins, interleukin-17 (IL-17) has been identified to play an important role in host defense against infections and influencing inflammatory responses particularly at the mucosal sites. This dual role makes IL-17 a molecule of interest, as it is involved in both host protection and pathogenesis of certain diseases [1]. The IL-17 cytokine family comprises six members that span from IL-17A to IL-17F, each differing in their amino acid composition. Notably, IL-17E is the most distinct, sharing only 16% similarity to IL-17A [2]. Among all the IL-17 members, IL-17A is the most thoroughly researched interleukin. IL-17 is secreted by various cell types, including T helper cell type 17 (T_h_17), gamma delta T cells, myeloid cells, and natural killer cells [3]. This broad range of cell sources underscores the versatile function of IL-17 in immune regulation and disease processes.

Interleukin-17 is recognized for its protective role against bacterial and fungal infections [4]. It promotes immunity against *K. pneumoniae*, *S. pneumoniae*, *M. tuberculosis* [5,6]. Its role in antifungal defence is highlighted by its ability to stimulate the production of pro-inflammatory cytokines, antifungal peptides and chemokines during candida albicans infections [7]. Additionally, patients treated with IL-17 inhibitors demonstrate an increased risk of oropharyngeal, esophageal, and cutaneous candidiasis [8].

Several studies indicated its pathogenic role against inflammatory, autoimmune diseases and viral diseases. An imbalance in IL-17 levels contributes to dysbiosis of the microbiome, thereby increasing the risk of developing autoimmune conditions [9]. Moreover elevated IL-17 levels have been associated with increased severity of coronavirus-related illness, including COVID-19 where excessive IL-17 secretion has been reported to worsen clinical symptoms [10]. Additionally, IL-17 expression is elevated in muscle, skeletal tissue, and serum of individuals affected with the Ross River virus (RRV). Inhibition of IL-17 has been shown to downregulate pro-inflammatory genes, thereby reducing tissue and cartilage damage in RRV-induced arthritis [11]. IL-17C has been shown to protect peripheral nervous system during HSV reactivation [12]. Additionally, IL-17 stimulates the secretion of other pro-inflammatory cytokines and promotes the accumulation of leukocytes, especially neutrophils, in patients suffering from Dengue infection [3]. Interestingly, increased IL-17 levels have been correlated with higher concentrations of Hepatitis C virus in the blood of infected patients, while Th17 cells from the same patient showed an inverse relationship thereby suggesting a complex interplay among IL-17 and Hepatitis C virus [13].

In addition to its role in virus biology, IL-17 is a key mediator in several inflammatory diseases. IL-17A predominantly affects epithelial cells, contributing to the development of psoriasis, a chronic inflammatory skin condition that can progress to psoriatic arthritis and atherosclerosis [14]. IL-17 also plays a role in inflammatory arthritis, where elevated levels in the bloodstream cause changes in blood vessels and liver cells, leading to higher production of c-reactive protein and weakening of muscle cells [15]. In the brain, IL-17 is crucial for brain immunity, with its deficiency linked to reduced short-term memory and synaptic plasticity [16]. Additionally, IL-17 offers potential therapeutic benefits in breast cancer by inhibiting metastasis as it polarizes neutrophils to supress cytotoxic T cells responsible for establishing metastasis [17].

Given the important role of IL-17 in regulating inflammatory responses, it is considered a promising target for drug development. Recent progress in therapeutics has focussed on identifying novel alternative therapeutic molecules, with particular emphasis on therapeutic peptides. Machine learning approaches have been successfully used to classify several therapeutic peptides [18–22]. Owing to the importance of IL-17 in the inflammatory pathways and disease states, we focussed on the classification of IL-17 inducing peptides. A literature review highlighted a 2017 study by Gupta et al. which aimed to distinguish IL-17-inducing peptides from non-inducing ones [23]. The study retrieved 338 IL-17 inducing and 984 non-inducing peptides from the Immune Epitope Database (IEDB) [24] that served as positive and negative datasets respectively. During the internal and external validation, MCC of 0.62 and 0.57 were achieved using the Support Vector Machine (SVM) classifier with dipeptide composition (DPC) of the peptide sequences. However, leveraging advancements in alignment-based approaches, machine learning techniques, and the availability of a larger dataset of experimentally validated IL-17-inducing and non-inducing peptides, we developed an improved three-stage hybrid method for classification of IL-17 inducing peptides. We retrieved 894 and 3124 IL-17 inducing and non-inducing peptides respectively from the IEDB database. We employed Basic Local Alignment Search Tool (BLAST), an alignment-based method to classify the peptides based on the similarity to known IL-17 and non-IL-17 peptides. MERCI was then used to identify the motifs that are exclusively present in IL-17 inducing peptides. We used the compositional module of the Pfeature algorithm to compute sequence-based features for each peptide sequence and developed machine learning models including k-Nearest Neighbour (kNN), Support Vector Machine (SVM) and Random Forest (RF) using these features. We found that while the model built via Support Vector Machine (SVM) using dipeptide composition (DPC) feature outperformed others, no single method was a clear winner. To address this, we designed a three-stage hybrid approach integrating BLAST, MERCI, and machine learning models (based on 400-DPC features), ensuring efficient classification of IL-17-inducing peptides from non-inducing ones. We report an accuracy of 88.08% and a Matthews Correlation Coefficient (MCC) of 0.68 on an external dataset surpassing these metrices of the existing methods. The models were subsequently incorporated into the webserver, providing the scientific community with a reliable tool to accurately predict the IL-17 induction potential of query peptide sequences. The overall experimental design is illustrated in Figure 1.

**Figure 1:**
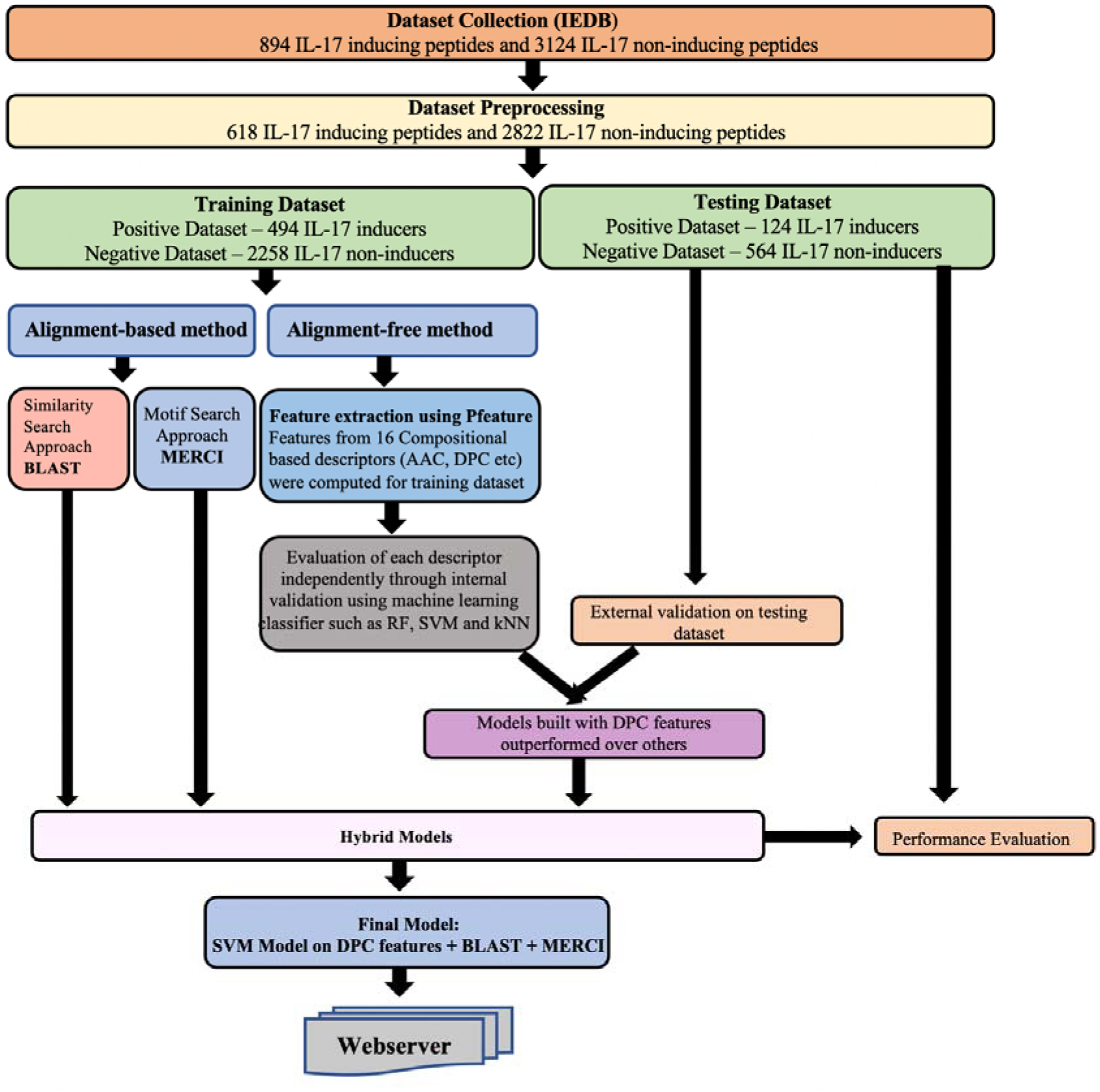
The overall design of IL17pred. The positive and negative peptides were obtained from the Immune Epitope Database (IEDB) database. These datasets were then split in 80:20 to form the training and the testing datasets respectively. The features of the peptides were computed using the Pfeature algorithm. A three-stage hybrid approach integrating BLAST, MERCI, and machine learning techniques was designed to achieve highly efficient classification of IL-17 inducing peptides. A user-friendly web server was also designed to enable the scientific community to predict the IL-17 inducing potential of the query peptide sequence.

## 2. Material and Methodology

### 2.1. Dataset Preparation

We retrieved the experimentally validated IL-17 inducing and non-inducing linear peptides from the Immune Epitope Database (IEDB) [25] on 27 January 2024. These peptides were either validated in human or mouse hosts. The positive and negative dataset consists of 894 IL-17 inducing and 3124 non-IL-17 inducing peptides, respectively. All the duplicate peptides as well as the peptides containing unnatural amino acids were removed from the study. The common peptides in both the positive and negative datasets were also eliminated. After that, we carried out a length-based analysis to find the distribution of peptide length among both datasets. We observed that the majority of peptides had a length between eight to thirty amino acids. We, therefore, removed those peptides that do not fall in this range from the datasets as the inclusion of these peptides may form an outlier and reduce the efficiency of prediction models [26]. Following this, the positive dataset included 618 IL-17 inducing peptides, while the negative dataset contained 2881 IL-17 non-inducing peptides. We employed CD-HIT-2D [27] with a 0.7 sequence identity threshold between the positive and negative datasets. This process identified and removed the peptide sequences from the negative dataset, that shared at least 70% similarity with those in the positive dataset. Eventually, the final dataset consists of 618 IL-17 inducing peptides as a positive dataset and 2822 non-IL-17 inducing peptides as a negative dataset.

### 2.2. Amino-acid and Dipeptide Compositional Analysis

To design IL-17 inducing peptides, it is critical to analyze the sequence of their constituent amino acids. We calculated the average composition of amino acids and dipeptides for both the IL-17 inducers and non-IL-17 inducers. Welch’s T-test was employed to identify the statistically significant amino acids and their dipeptides enriched in IL-17 inducing peptides compared to non-inducing peptides and vice versa.

### 2.3. Positional Preferences of Amino-acids

The analysis of amino acid composition indicated that certain amino acids were significantly abundant in IL-17 inducers as compared to the non-IL-17 inducers. To visualize the distribution of these enriched amino acids, we created a Two Sample Logo (TSL) [28]. Since the TSL tool requires peptides of a fixed length, we followed a commonly used protocol to create the peptides of a fixed length [29,30]. Given that the minimum length of the peptides in our dataset was eight amino acid residues, we extracted the first eight amino-acids from the N-terminus of the peptide and the last eight amino-acids from the C-terminus of the peptide, combining them to form a 16 amino-acid long peptide. This process was repeated for all the peptides from the positive and negative datasets.

### 2.4 Alignment-based Approach

#### 2.4.1. BLAST: Similarity Search Method

Basic Local Alignment Search Tool (BLAST) is a widely used method for annotating peptide and nucleotide sequences based on the similarity search [31]. We implemented blastp-short (protein–protein BLAST) suite of BLAST+ version 2.6.0 to develop the similarity-based search module [32]. Here, the query protein sequence was hit against the database of known IL-17 and non-IL-17 inducers. We adopted the two strategies i.e. top-hit and voting strategy, widely used in literature to annotate the query peptide sequence [33]. The voting strategy is an ensemble of top 5 hits. If the query sequence gets less than 5 hits against the database, then the query sequence was annotated as IL-17 inducer or non-IL-17 inducers based on the first hit. However, if the number of hits for a query sequence becomes more than 4, then the query sequence is assigned as IL-17 or non-IL-17 inducers on the basis of maximum hits. The sequence is annotated as IL-17 inducer if the top five hits have maximum IL-17 inducers. Similarly, if the top five hits have maximum non-IL-17 inducers, then the query sequences are classified as non-IL-17 inducers. The performance of the method was assessed based on the various E-value cut-offs.

#### 2.4.2. MERCI: Motif Search Method

The identification of motifs in peptide sequences is essential for determining their functional annotations. We utilized the freely available software Motif-EmeRging and Classes-Identification (MERCI) [34], motifs within IL-17 and non-IL-17 inducing peptides. This software was implemented to extract recurring patterns from both positive and negative datasets using the default parameters.

### 2.5. Alignment-free Approach using Machine Learning

#### 2.5.1 Feature Extraction

To develop any machine learning based prediction model in the biological sciences, sequences must be represented as numerical vectors, which are then utilized by the models for further analysis. The Pfeature algorithm [35] is one of the feature extraction methods employed to derive features from the peptide sequences. The features computed by the Pfeature algorithm are widely used to build classification models [29,32,36]. We used the 16 descriptors of the compositional module of the Pfeature algorithm to extract features of peptide sequences for both positive and negative datasets. The descriptors and their corresponding vector lengths are tabulated in supplementary table 1. By combining the features from each descriptor, we computed a total of 9151 features for each peptide sequence.

#### 2.5.2. Machine Learning Classifiers

The machine learning techniques commonly applied in biological classification problems including k-Nearest Neighbour (kNN), Random Forest (RF), and Support Vector Machine (SVM) were used to generate robust and generalized prediction models. These techniques were implemented using the scikit package from Python [37]. Hyperparameter tuning was conducted through grid search cross-validation and the performance metrics were reported based on the optimal parameters.

##### 2.5.2.1. *k*-Nearest Neighbour (kNN)

The *k*-Nearest Neighbour (kNN) algorithm is a versatile and intuitive classification method widely used to build decision model [38]. kNN is non-parametric in nature. It operates on the principle of proximity, where a data point is classified based on the class labels of its nearest neighbours in the feature space. kNN does not assume any underlying probability distribution of the data and is particularly effective in handling non-linear class distributions.

##### 2.5.2.2. Random Forest (RF)

The Random Forest (RF) algorithm is an ensemble learning method composed of multiple decision trees. Each decision tree in the random forest is built using a different subset of the training data, leading to diverse individual trees with unique predictive capabilities [39]. Through a process called bagging (bootstrap aggregating), where each tree is trained on a bootstrapped sample of the data, random forest combines the predictions of these individual trees to make a final decision. By aggregating the decisions of multiple trees, random forest can mitigate overfitting and variance, leading to improved generalization performance compared to single decision tree models. This collective decision-making process enhances the robustness and reliability of the random forest algorithm.

##### 2.5.2.3. Support Vector Machine

The Support Vector Machine (SVM) [40] is a powerful algorithm employed for classification problems. It determines an optimal decision boundary by maximizing the margin between hyperplanes that pass through the support vectors of different classes. These support vectors are crucial data points that influence the determination of the decision boundary. SVM is renowned for its ability to handle both linear and non-linear classification problems efficiently. It achieves this by utilizing various kernel functions such as linear, polynomial, radial basis function (RBF), and to transform the input space into a higher-dimensional space where linear separation becomes possible. This capability allows SVM to effectively model complex relationships within data, making it a versatile tool in classification tasks across various domains.

#### 2.5.3 Internal and External Validation

The positive and negative dataset comprise 618 IL-17 inducing and 2822 non-IL-17 inducing peptides respectively. Following the standard protocol, we divided the dataset into training data (80%) and testing data (20%). The training dataset includes 494 IL-17 inducing peptides and 2258 non IL-17 inducing peptides. The testing dataset consists of 124 known IL-17 inducers and 564 non IL-17 inducing peptides. The training dataset was used to train the models while the testing dataset was employed to assess the model’s performance. Internal validation was employed to train, test and evaluate the prediction models. During 5-fold cross validation, the training dataset was split into 5 equal parts. Each of the five parts consists of an equal number of positive and negative peptide sequences. The models were trained on four parts while the fifth part was used for testing. This process was iterated five times so that each part could act as testing data. The performance metrics from each iteration in the 5-fold cross-validation were averaged to produce the final metrics. We performed external validation to evaluate the efficiency of the trained model.

#### 2.5.4 Evaluation Parameters

In a binary classification problem, the data must be categorized into the positive and negative classes. Accordingly, peptides that have been experimentally validated and shown to induce IL-17 were designated as the positive class (P), while peptides incapable of inducing IL-17 were designated as the negative class (N). The performance of prediction models was assessed using standard evaluation metrics encompassing both threshold dependent and independent parameters. The sensitivity, specificity, accuracy and Mathews Correlation Coefficient (MCC) are commonly used threshold dependent parameters. Sensitivity, also known as True Positive Rate (TPR), is the ratio of correctly predicted positive labels to the total positives. Specificity is the ratio of correctly predicted negative labels to the total negatives, also known as the true negative rate (TNR), Accuracy is the ratio of total correct prediction labels (both true positive and true negatives) to the total number of labels. MCC is a statistical metric for handling class-imbalance problems. MCC yields a high score when the model predicts both the positive and negative labels correctly. These parameters were calculated using the following equations

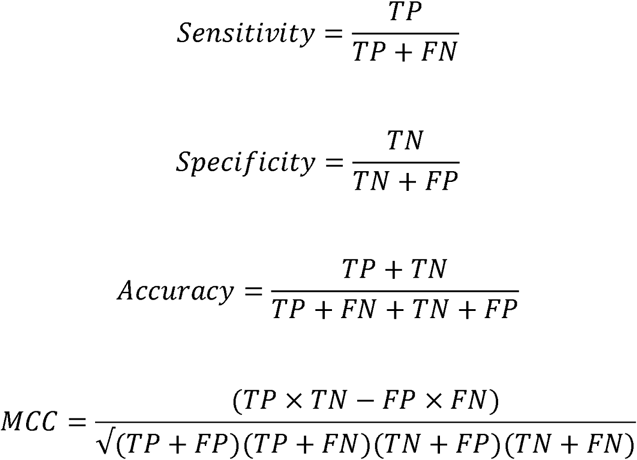

where TP refers to number of true positives labels; TN refers to numbers of true negatives labels; FP, refers to False Positives, is the number of negative labels classified as positive labels; FN refers to False Negatives, is the number of positive labels classified as negative labels.

In addition to the above-mentioned threshold dependent parameters, threshold independent parameter is extensively used in many studies for evaluating the performance of machine learning prediction models and is well established in the literature [33,36]. The area under the receiver operating characteristics curve is one such threshold independent parameter which is calculated via the plot between the Sensitivity (True Positive Rate) and 1-Specificity (False Positive Rate).

### 2.6. Proposed Hybrid Approach

Hybrid approaches combining multiple prediction algorithms have recently been employed for biomolecule classification [32,33,41]. These methods utilize various algorithms to assign scores to potential positive and negative peptides ultimately facilitating accurate biomolecule prediction. In this study, a three-stage hybrid approach was applied integrating algorithms at different stages, including BLAST, motif search and an alignment-free machine learning method, to assess the IL-17 induction potential of the query peptides. In the first stage, peptide sequence alignment was performed using the BLAST algorithm predicting peptide sequences at an E-value threshold of 10 . Scores were assigned based on prediction outcomes: a correct positive prediction received a score of ‘+0.5’, a correct negative prediction was ‘-0.5’, and no hits were assigned a score of ‘0’.

In the second stage, the same peptide sequences were analysed using the MERCI algorithm. Similar scoring criteria were applied: ’+0.5’ if a motif was identified in the positive dataset, ’-0.5’ if a motif was found in the negative dataset, and ’0’ if no motif was identified. In the final stage, machine learning-based methods analysed the peptides. The scores from BLAST and MERCI were combined with the machine learning results to predict the IL-17 induction potential of the peptides.

### 2.7. Design of Web-based Prediction Tool

To facilitate the scientific community, we designed and developed a user-friendly web server, iIL17pred (http://www.soodlab.com/iil17pred/), to classify IL-17 inducers from non-IL-17 inducers. Ubuntu was used to host the site on an Amazon Web Services (AWS) LightSail server instance. The front end was built using HTML, CSS, and JavaScript to structure the pages, add design elements, and provide interactivity such as form validations and drop-down menus. A Python program within the Flask application, a small and lightweight Python web framework that provides useful features for web development, facilitated communication with the machine-learning model, delivering predictions to the web application’s front end. The users are allowed to input a peptide sequence, which is then processed and provides the probability score indicating the IL-17 induction potential of the query peptide. Besides the predict module, the webserver has design, protein scan, blast and motif search functionalities. The design module in iIL17pred facilitates the generation of single amino acid mutants of the given peptide sequence. The module then predicts the IL-17 induction potential of all mutants. This enables the users to design novel IL-17 inducing peptides. The protein scan module generates and predicts the IL-17 induction potential of all possible overlapping peptides from a protein sequence. This module allows the user to extract the IL-17 inducing peptide from a protein sequence. The blast module in iIL17pred helps the user to classify the peptide sequences based on their similarity to known IL-17 inducing peptides. This module predicts the sequence as an IL-17 inducer if it gets hit from the database. The motif search module facilitates the user to find the IL-17 inducing motifs in the query sequence.

## 3. Results

### 3.1. Compositional Analysis of IL-17 and Non-IL-17 Inducing Peptides

We computed the average amino acid and dipeptide composition in IL-17 inducing and non-inducing peptides. Welch’s T-test was used to compute the statistically significant enriched amino acids and dipeptides among both groups. Figure 2A shows the average amino acid composition of 20 amino acids. It was observed that some amino acids including Leucine (L), Arginine (R), Serine (S), and Tryptophan (W) are significantly enriched in IL-17 inducing peptides (Welch’s *t*-test, *p* < 0.01) whereas amino acids like Aspartic acid (D), Glutamic acid (E), Glycine (G), Lysine (K) and Valine (V) are significantly abundant in non-IL17-inducing peptides (Welch’s *t*-test, *p* <0.01) (Supplementary Table 2). The analysis of dipeptide composition reveals that some dipeptides were significantly enriched (Welch’s *t*-test, *p* < 0.01) in IL-17 inducing peptides whereas some were enriched in the non-IL-17 inducing peptides (Supplementary Table 3). Figure 2B represents the top 15 statistically significant enriched dipeptides in IL-17 inducers and includes AL, LK, PL, LS, IL, LR, IS, SV, PS, RL, SF, RS, YF, RP and FF whereas dipeptides including VV, DV, VD, VA, AG, DA, DK, EE, VK, ED, ID, KE, EA, RG and AD were statistically significantly abundant in non-IL-17 inducers. The data suggests that these dipeptide signatures could be used as the distinguishing features among the positive and negative sets of the peptides.

**Figure 2:**
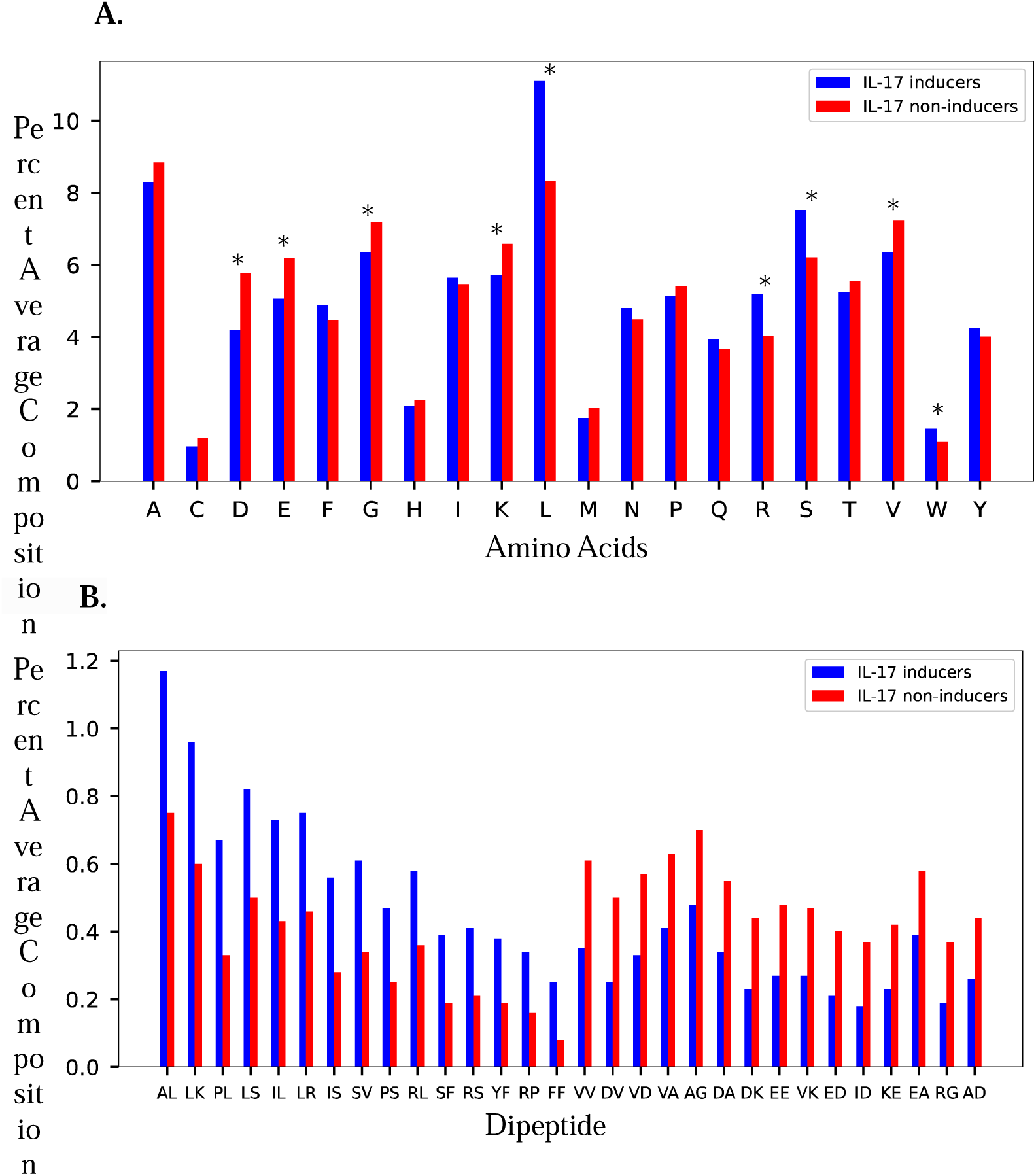
Comparative analysis of the average amino acid and dipeptide composition among the IL-17 and non-IL-17 inducers. (A) The amino acids are represented by the X-axis whereas the Y-axis represents the average composition of amino acids in percentage. The statistically significant enriched amino acids (p-value < 0.01) in IL-17 inducers and IL-17 non-inducers are represented by *. (B) The X-axis represents the dipeptides whereas the Y-axis represents the average composition of dipeptides in percentage. The top 15 statistically significant enriched dipeptides (p-value < 0.01) in IL-17 and non-IL-17 inducers are shown.

### 3.2. Positional Analysis of IL-17 Inducing and Non-IL-17 Inducing Peptides

Compositional analysis of the IL-17 inducing peptides revealed that some amino acids were highly enriched in the positive dataset as compared to the negative one. To identify the position of these enriched amino acids, we created the Two-sample logo (TSL) plot for both the IL-17 inducing (positive) and non-inducing (negative) peptides (Figure 3). As mentioned above, in the TSL plot, the amino acids from positions 1-8 represent the first eight N-terminus amino acids of a peptide while the amino acids from 9-16 represent the last eight C-terminus amino acids of a peptide. We observed that leucine was preferred at both the N- and C-terminus of IL-17 inducing peptides. It was mostly preferred at 2^nd^, 5^th^, 7^th^, 10^th^, 12^th^, 13^th^, 14^th^ and 16^th^ position in a peptide sequence. In addition, the amino acids ‘Y’, ‘R’, ‘S’, ‘F’ and ‘W’ were highly preferred at most of the positions in a positive dataset. On the contrary, the amino acid ‘D’, ‘E’ and ‘K’ were highly preferred at most of the positions in IL-17 non-inducing peptides. The data suggested that IL-17 inducing peptides might consist of some signature amino acids at specific positions and thus could act as distinguishing features in the classification of peptides.

**Figure 3:**
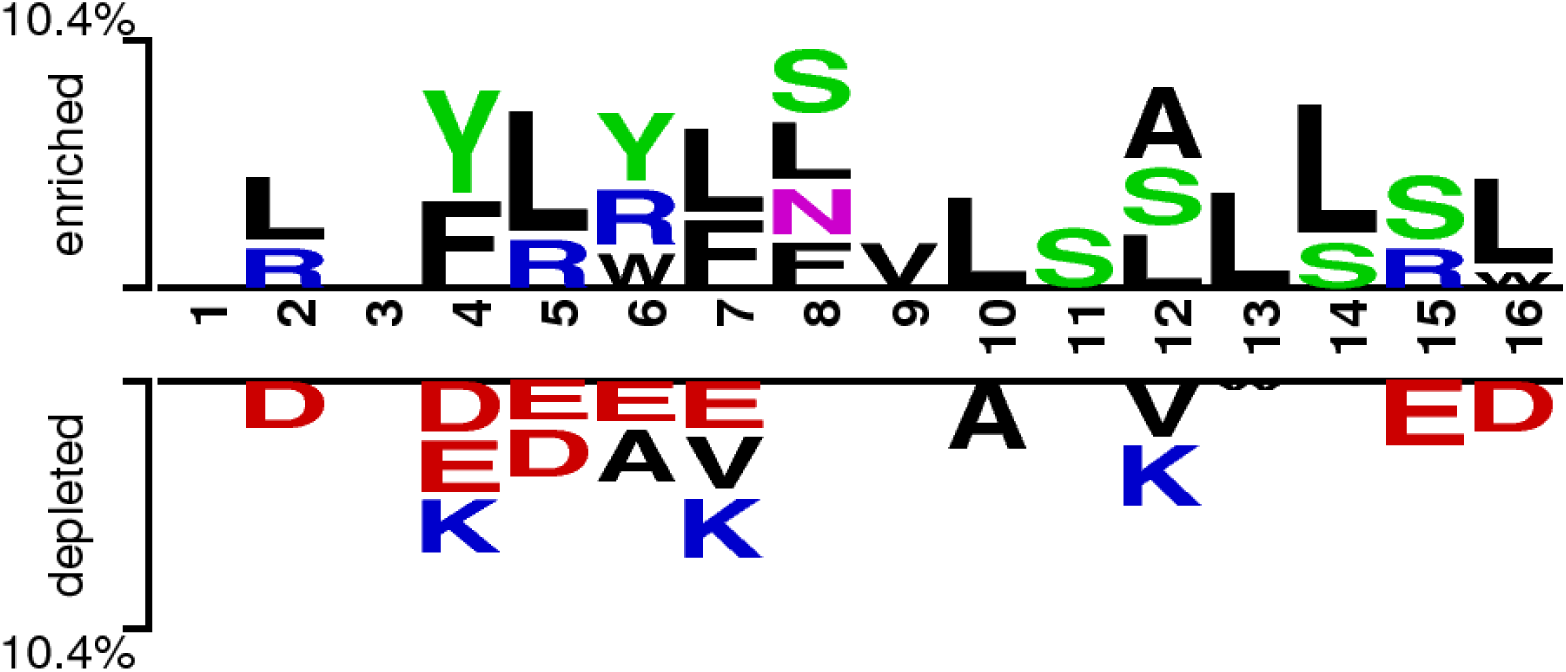
The Two-sample Logo (TSL) plot of IL-17 inducing and non-IL-17 inducing peptides illustrating the distribution of enriched amino acids in both datasets.

### 3.3. Alignment-based Methods

#### 3.3.1. BLAST – Similarity Search Method

BLAST is used to annotate the function of unknown peptide or protein sequence based on similarity searches against a database of known peptides. In this study, we leveraged this tool for developing alignment-based models to classify the IL-17 inducers from non-IL-17 inducing peptides. Since the constituent peptides in our datasets (both positive and negative) range from 8-30 amino acids, hence, we implemented blastp-short (protein–protein BLAST) suite across the e-values 10^-6^ to 10^-1^. To evaluate the efficacy of alignment-based model and avoid any biases we performed five-fold cross-validation. In this, we split the training dataset into 5 equal folds, where the peptide sequences in the 4 folds were used to construct the BLAST database, and the sequences in the remaining fold were searched against the above database for annotation. The entire process was iterated 5 times. To evaluate the performance of the alignment-based model on the validation dataset, we build the database from the entire training dataset. This database is used to annotate each peptide in the validation dataset using the similarity search method. In our study, for the training dataset, the number of correct hits (sensitivity) rises from 0.69% to 4.82% whereas for the validation datasets, the number of correct hits rises from 0.29% to 5.23%. In addition to this, for the training and validation dataset, we attained specificity ranges from 1.96% to 57.46% and 0.43% to 62.79% respectively across E-value ranges from 10^-6^ to 10^-1^. We can infer from table 1 that for both the training and validation datasets, the number of incorrect hits, or errors, rises proportionately to the number of correct hits indicating that BLAST approach suffered from a high error rate and was thus not sufficient alone to classify IL-17 inducing peptides from non-IL-17 inducers.

**Table 1:** The performance of BLAST-based approach on the positive and negative dataset across different e-values.

| E-value | Training |  |  |  | Validation |  |  |  |
| --- | --- | --- | --- | --- | --- | --- | --- | --- |
|  | IL-17 inducers |  | IL-17 non-inducers |  | IL-17 inducers |  | IL-17 non-inducers |  |
|  | Correct hits (%) | Wrong hits (%) | Correct hits (%) | Wrong hits (%) | Correct hits (%) | Wrong hits (%) | Correct hits (%) | Wrong hits (%) |
| $10^{-1}$ | 133 (4.82%) | 28 (1.01%) | 1583 (57.46%) | 25 (0.91%) | 36 (5.23%) | 1 (0.15%) | 432 (62.79%) | 9 (1.31%) |
| $10^{-2}$ | 118 (4.28%) | 16 (0.58%) | 1184 (42.98%) | 16 (0.58%) | 29 (4.22%) | 0 (0%) | 324 (47.09%) | 7 (1.02%) |
| $10^{-3}$ | 76 (2.76%) | 7 (0.25%) | 489 (17.75%) | 9 (0.33%) | 16 (2.33%) | 0 (0%) | 130 (18.89%) | 6 (0.87%) |
| $10^{-4}$ | 46 (1.67%) | 3 (0.11%) | 266 (9.66%) | 7 (0.25%) | 10 (1.45%) | 0 (0%) | 77 (11.19%) | 5 (0.73%) |
| $10^{-5}$ | 30 (1.09%) | 2 (0.07%) | 108 (3.92%) | 4 (0.15%) | 4 (0.58%) | 0 (0%) | 29 (4.22%) | 4 (0.58%) |
| **10<sup>-6</sup>** | 19 (0.69%) | 0 (0%) | 54<br>(1.96%) | 1<br>(0.04%) | 2<br>(0.29%) | 0 (0%) | 3 (0.43%) | 3 (0.44%) |

#### 3.3.2 MERCI – Motif Search Method

Motifs are the short patterns within the peptide sequences that are responsible for their functions. An insight into the motifs of a peptide sequence helps in predicting the function of the peptide. Among various tools for the motif prediction, MERCI has been widely used to identify motifs among a set of peptides [20,41]. In this study, we used MERCI algorithm to identify the molecular determinants that are exclusively present in positive and negative datasets. Analysis of the peptide sequences revealed that some motifs including PVI, FSR, FSRV and NPVI, were exclusively present in the IL-17 inducing training peptides, whereas some motifs, including GPA, AGP and DVD were present in the non-IL-17 inducing peptides. Additionally, several other motifs in the positive and negative datasets were identified which are described in the supplementary table 4. Identification of different motifs in the positive and the negative datasets can potentially lead us towards the classification of the peptides and hence this data was used successfully towards the classification of the IL-17 inducing peptides.

### 3.4. Machine Learning Prediction Models

We employed the Pfeature algorithm to compute the 9151 features of IL-17 and non-IL-17 inducing peptides. We computed 16 compositional based descriptor such as AAC, DPC, TPC etc. and built machine learning models with each descriptor to delineate their roles in distinguishing IL-17 from non-IL-17 inducers. In addition to these, 9151 features were used to build the machine learning based prediction model by using three different machine learning classifiers i.e. *k*-Nearest Neighbour (kNN), Support Vector Machine (SVM), and Random Forest (RF). To optimize the machine learning models, we performed hyperparameter tuning via grid search. To evaluate the efficacy of each model, we performed stratified 5-fold cross validation on the training dataset and calculated average sensitivity, specificity, accuracy and MCC. Further, we also assessed the performance of our model on the testing set that was never used at any stage of model building. The performance of SVM based models on different compositional features are discussed in table 2. The effectiveness of kNN and RF based models on each compositional feature are discussed in supplementary tables 5 and 6 respectively. Figure 4 and Supplementary Figure 1 and 2 demonstrates that the machine learning based prediction models built using dipeptide composition (DPC) features exhibited superior performance compared to other feature types. The model built using DPC features in conjunction with SVM, kNN and RF achieved an MCC of 0.63, 0.56 and 0.55 on external validation data respectively. The model built on DPC features has shown superior performance in classifying biomolecules in several other studies [23,32]. Figure 5, shows the model built using SVM classifier with DPC features surpassed the performance of other classifiers with an accuracy and MCC of 84.57%, and 0.6 respectively on internal validation while achieving an accuracy and MCC of 85.76% and 0.63 on external validation.

**Table 2:**
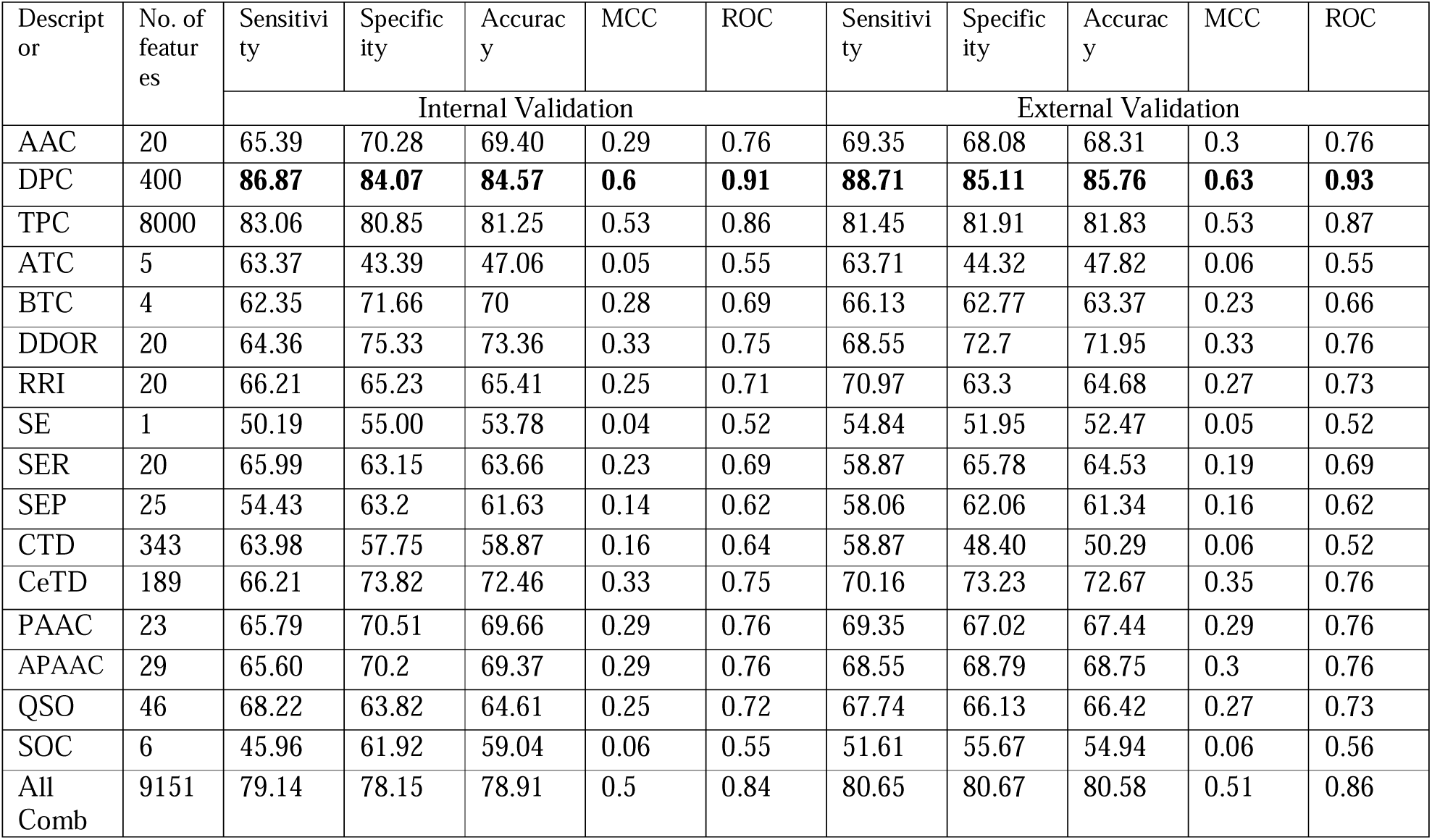
Performance-metrics of machine learning based models on internal and external validation developed by utilizing various compositional features in conjunction with Support Vector Machine (SVM) classifier. The maximum values in each performance metrics are highlighted in bold. Abbreviation: AAC, Amino Acid Composition; DPC, Dipeptide Composition; TPC, Tripeptide Composition; ATC, Atomic Composition; BTC, Bond Composition; DDOR, Distance distribution of residue; RRI, Residue repeat Information; SE, Shannon-entropy of protein; SER, Shannon entropy of all amino acids; SEP, Shannon-entropy of physiochemical property; CTD, Conjoint triad calculation of the descriptor; CeTD, Composition-enhanced transition distribution; PAAC, Pseudo amino acid composition; APAAC, Amphiphilic pseudo amino acid composition; QSO, Quasi-sequence order; SOCN, Sequence order coupling number; MCC, Matthews correlation coefficient.

**Figure 4:**
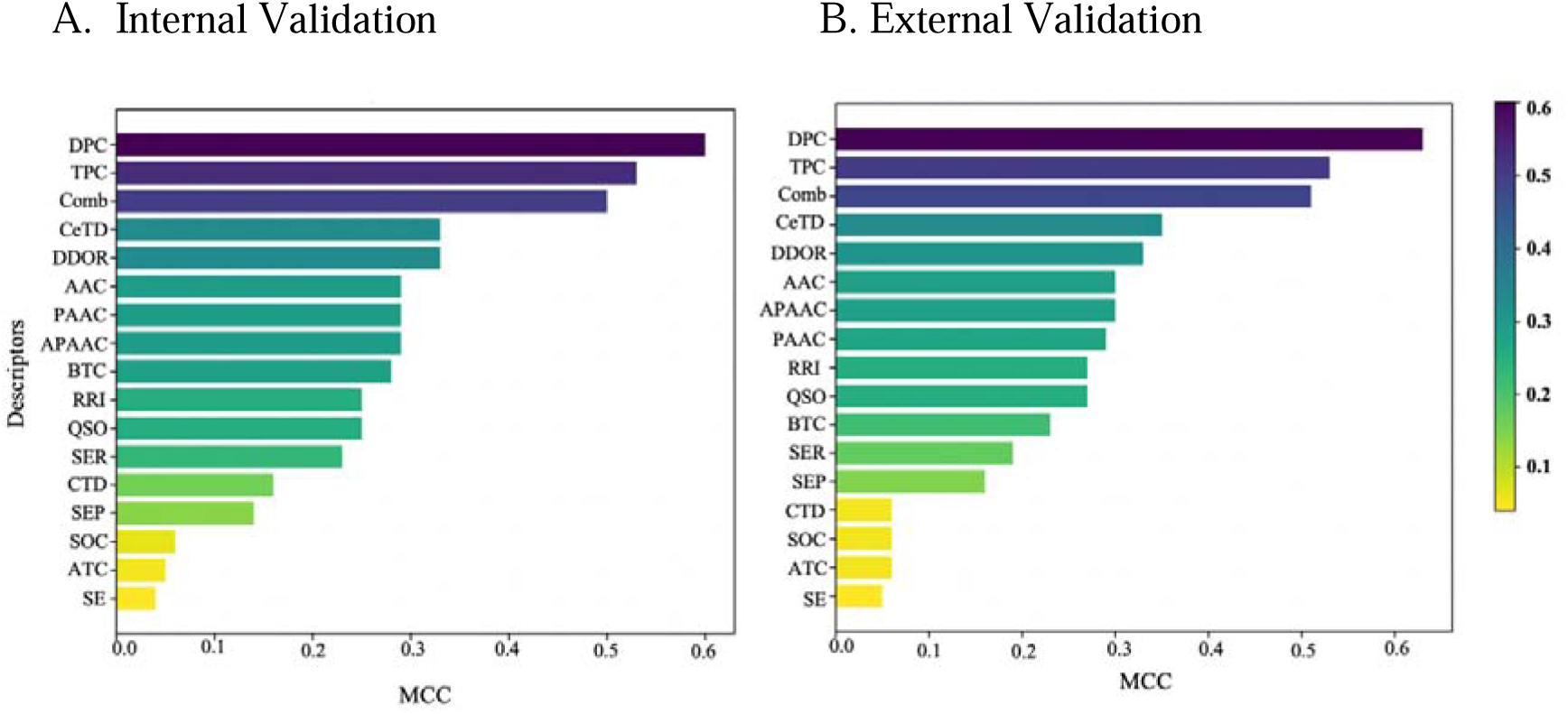
The representation of performance metrics of models using different compositional features with a Support Vector Machine (SVM) classifier. Matthews correlation coefficient (MCC) values are on the x-axis and the features are on the y-axis. The descriptors are ranked based on their MCC values, represented by color gradients. Higher MCC values indicate better performance in predicting outcomes.

**Figure 5:**
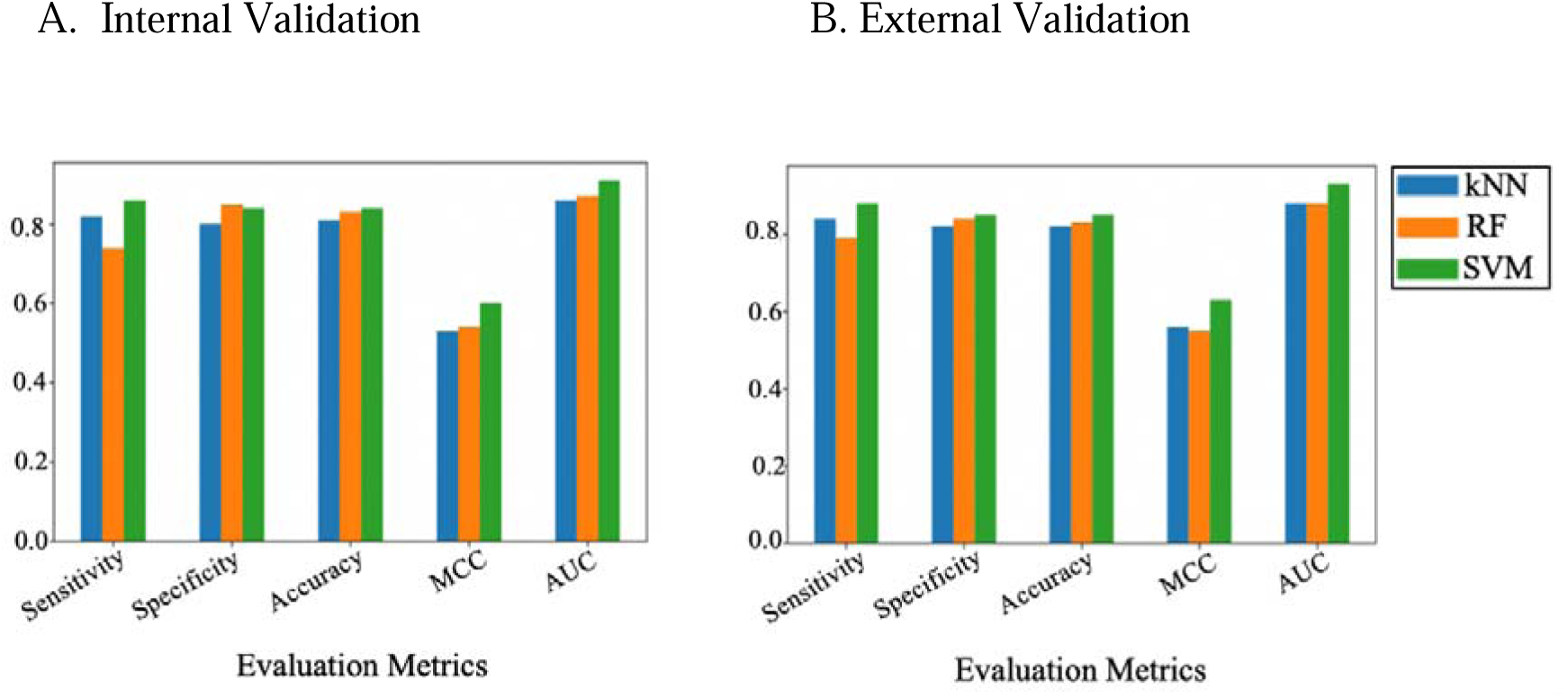
shows the performance of models built on dipeptide composition (DPC) features using different classifiers i.e. k-nearest neighbour (kNN), Random Forest (RF) and Support Vector Machine (SVM). The X-axis represents the evaluation metrics including sensitivity, specificity, accuracy, Matthews Correlation Coefficient (MCC) and Area under the curve (AUC) and the Y-axis represent the values.

### 3.5. ML-Based Models with BLAST Search

We observed from Figure 4 and Supplementary Figure 1 and 2, the model built on DPC features outperformed over other models. Hence, we combined the ML prediction model with alignment-based approach to efficiently classify IL-17 inducers from non-IL-17 inducers. we integrated the strengths of the alignment based method i.e. BLAST and an alignment-free method i.e. ML-based models. Initially, an alignment based search was performed on the query peptide sequence using BLAST. The query peptide was classified as IL-17 inducer or non-IL-17 inducing based on the hit against the database. If the query peptide sequence did not get any hit, then the dipeptide composition-based model is used to classify the respective sequence. [32,41]. The integration of BLAST with machine learning based prediction models leads to the efficient classification of IL-17 inducers from non-IL-17 inducers. The performance of BLAST combined with machine learning-based model on DPC features were discussed in table 3. From figure 6 and supplementary figure 3 and 4, it was observed that after integrating BLAST with ML models resulted in an increase in both accuracy and MCC. For instance, the SVM-based model showed an improvement in its performance metrics: the MCC increased from 0.60 to 0.64 on the training dataset and from 0.63 to 0.66 on the validation dataset. This demonstrates that combining sequence alignment methods with machine learning models can enhance the classification of IL-17 inducers and non-inducers.

**Table 3:** The performance of alignment based approach (BLAST) combined with machine learning-based model using DPC features. The maximum values in each performance metrics are highlighted in bold.

| Classifier | Sensitivity | Specifi<br>city | Accuracy | MCC | ROC | Sensitivity | Specifi<br>city | Accura<br>cy | MCC | ROC |
| --- | --- | --- | --- | --- | --- | --- | --- | --- | --- | --- |
|  | Internal Validation |  |  |  |  | External validation |  |  |  |  |
| SVM | **86.87** | **86.73** | **86.75** | **0.64** | **0.93** | **88.71** | **87.23** | **87.5** | **0.66** | **0.94** |
| kNN | 84.85 | 83.40 | 83.66 | 0.58 | 0.88 | 86.29 | 84.75 | 85.02 | 0.61 | 0.9 |
| RF | 79.8 | 86.28 | 85.11 | 0.58 | 0.88 | 83.06 | 85.63 | 85.17 | 0.6 | 0.9 |

**Figure 6:**
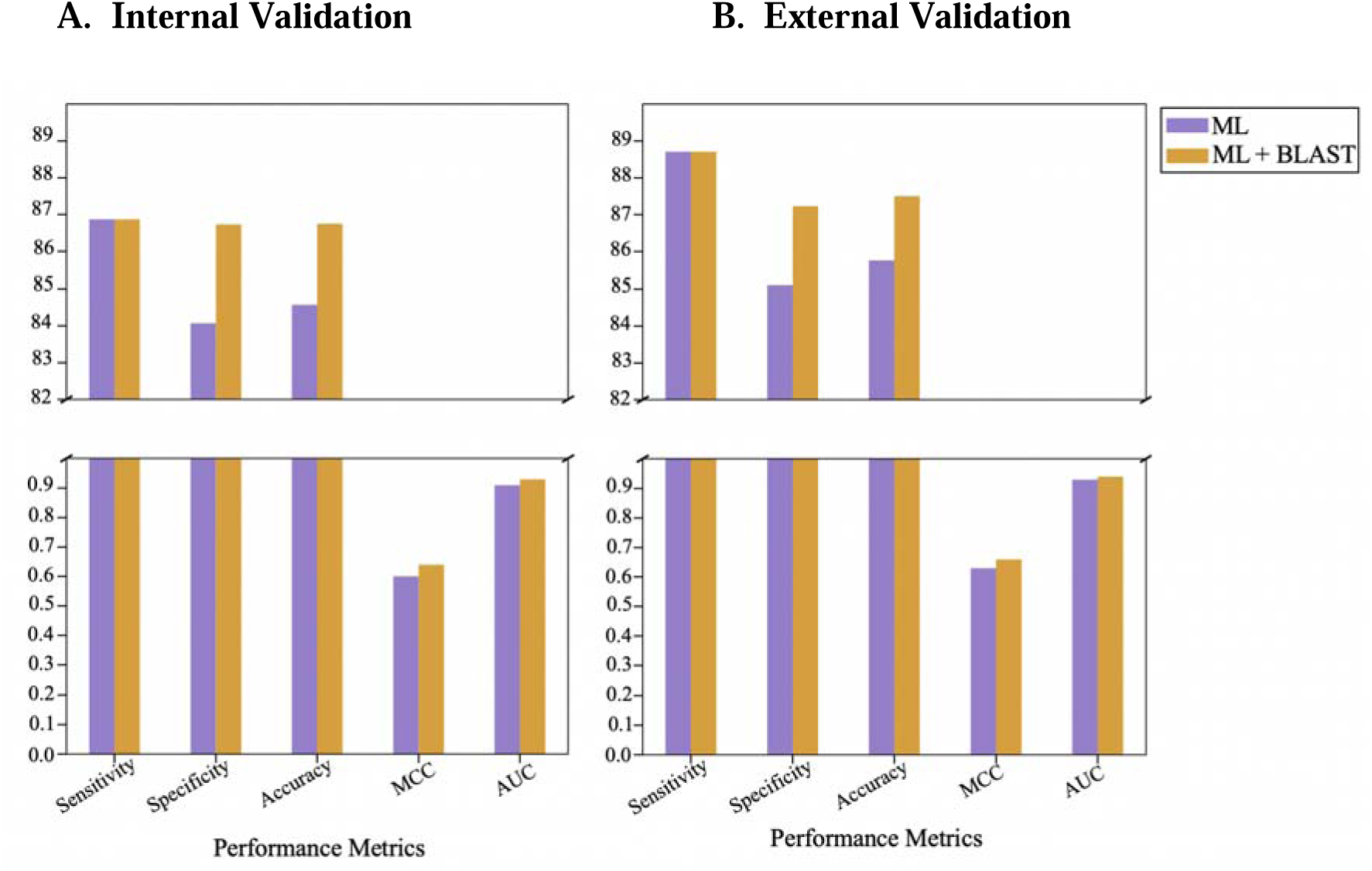
shows the comparison of performance of Support Vector Machine (SVM) based model alone and in combination with BLAST. These models were built on dipeptide composition (DPC) features. The X axis represent the performance metrics and Y axis represent the values.

### 3.6. ML-based Models with Motif Approach

Motifs are short sequences within peptides, that play a crucial role in determining their function. The MERCI algorithm was used to identify the motifs in training dataset. We leveraged the combined strengths of motif search method i.e. MERCI and ML-based models on DPC features. The given peptide was classified as either an IL-17 inducer or a non-IL-17 inducer based on the presence of specific motifs. If the motif corresponds to an IL-17 inducer, the peptide is classified as IL-17 inducing; if the motif corresponds to a non-IL-17 inducer, it is classified as non-IL-17 inducing. If the query peptide sequence did not match any known motifs, a dipeptide composition-based model is used to classify the sequence. The performance metrics discussed in table 4, and found that the SVM-based model with MERCI outperformed over other models and attained the MCC of 0.61 and 0.64 on training and validation dataset respectively. As observed from the figure 7 and supplementary figure 5 and 6, there is only a marginal improvement in performance metrics compared to when MERCI was not incorporated into the classification process.

**Table 4:** The performance of motif based approach (MERCI) combined with machine learning-based model using DPC features. The maximum values in each performance metrics are highlighted in bold.

| Classifier | Sensitivity | Specifi<br>city | Accuracy | MCC | ROC | Sensitivity | Specifi<br>city | Accura<br>cy | MCC | ROC |
| --- | --- | --- | --- | --- | --- | --- | --- | --- | --- | --- |
|  | Internal Validation |  |  |  |  | External validation |  |  |  |  |
| SVM | **87.88** | 84.13 | **84.75** | **0.61** | **0.91** | **89.43** | **85.66** | **86.34** | **0.64** | **0.93** |
| kNN | 83.84 | 82.52 | 82.76 | 0.56 | 0.88 | 85.37 | 82.65 | 83.14 | 0.57 | 0.88 |
| RF | 75.76 | **86.28** | 84.39 | 0.55 | 0.87 | 81.30 | 84.96 | 84.30 | 0.57 | 0.88 |

**Figure 7:**
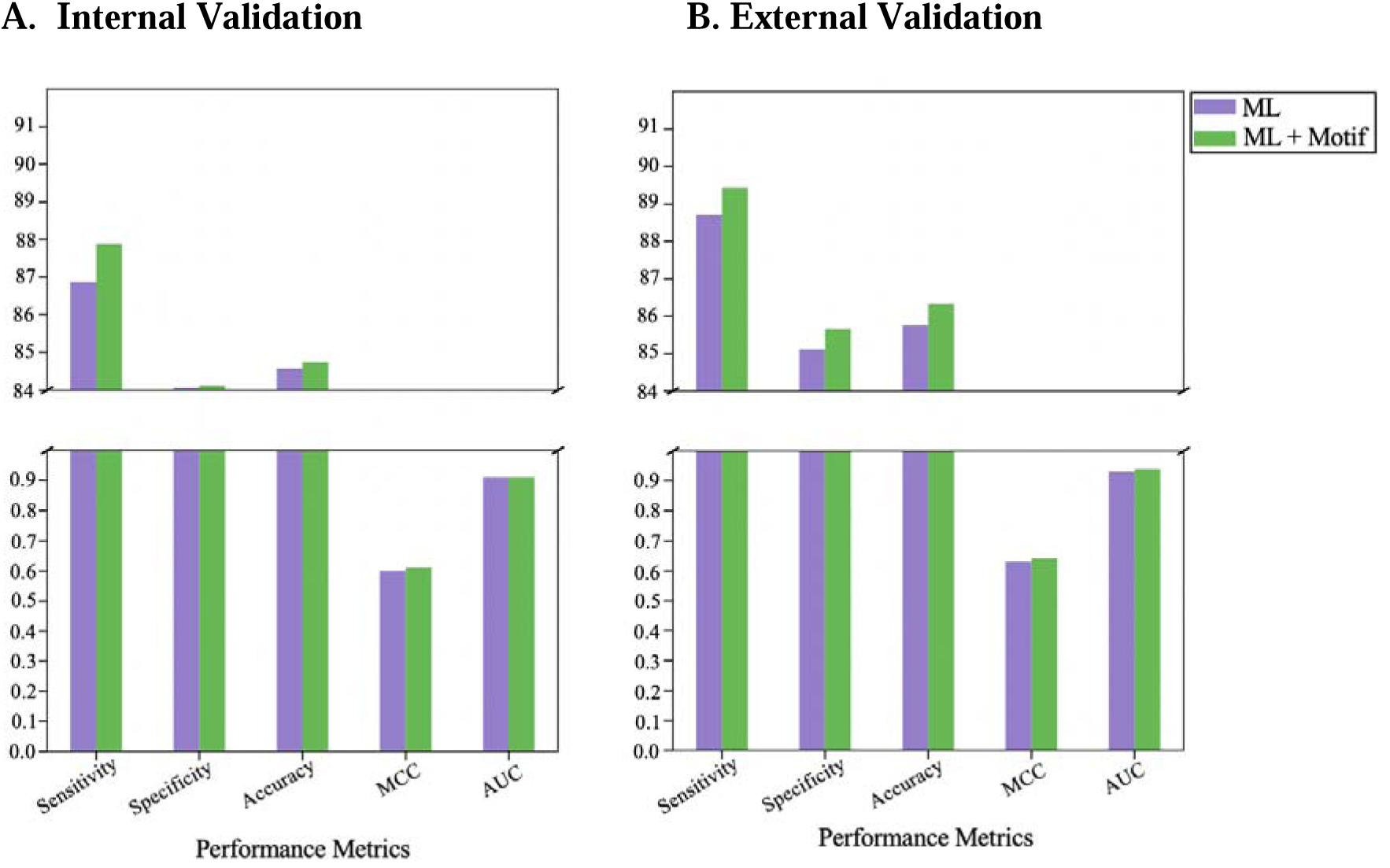
shows the comparison of performance of SVM based model alone and in combination with MERCI. These models were built on DPC features. The X axis represent the performance metrics and Y axis represent the values.

### 3.7 Proposed Hybrid Approach

Efficient classification of biomolecules is crucial in biology, as it provides valuable insights into their potential roles and functions. Accurate classification allows researchers to better understand molecular mechanism, predict biological activities, and develop targeted therapeutic interventions. The classification of the biomolecules can be achieved either through alignment-free method or alignment based method. However each method have its pros and cons. To overcome the drawbacks of individual methods, we implemented and explored the power of the hybrid approach that combine and leverages the strength of each individual approach i.e. BLAST, MERCI and ML. Initially, the query peptide sequences was classified by BLAST at an E-value of 10^-1^, followed by MERCI and then predicted by machine learning based prediction model that were trained on the DPC features. From figure 8 and supplementary figure 7 and 8, we observed a significant improvement in the accuracy and MCC during internal and external validation. As show in table 5, the SVM-based hybrid model surpassed other models and achieved an MCC of 0.65 and 0.68 during internal validation and external validation respectively. These results are consistent with earlier studies [32,33,41], demonstrating that a hybrid approach enhances the efficiency of biomolecule classification compared to individual methods.

**Figure 8:**
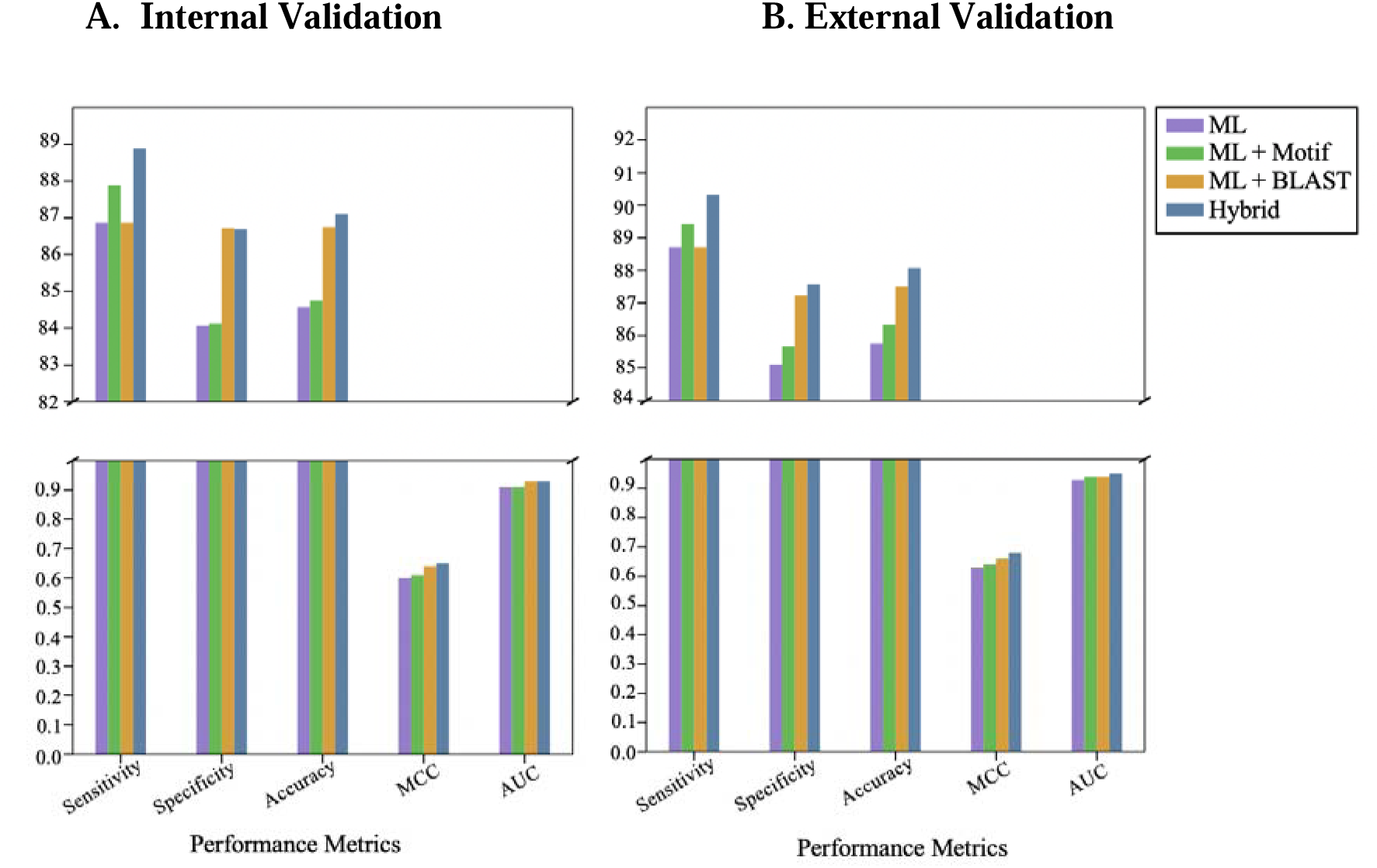
shows the comparison of performance of Support Vector Machine (SVM) based model and in combination with MERCI and BLAST, and hybrid model. These models were built on DPC features. The X axis represent the performance metrics and Y axis represent the values.

**Table 5:**
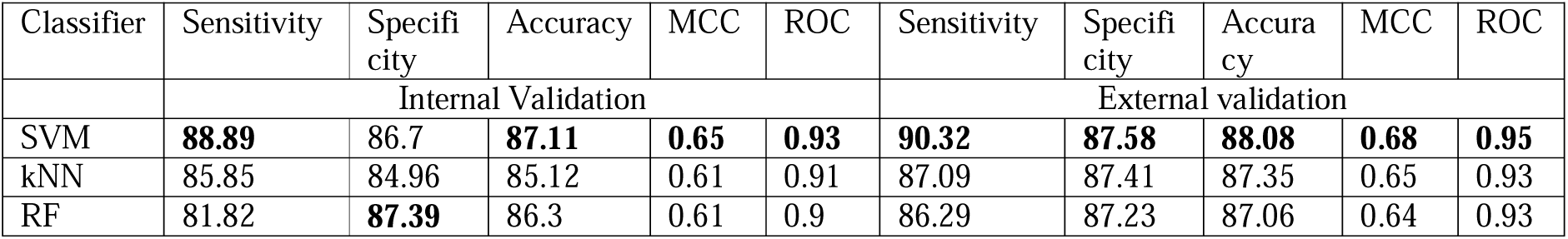
The performance hybrid approach combines ML model with alignment-based search using BLAST and motif-based search using MERCI. The maximum values in each performance metrics are highlighted in bold.

## 4. Discussion

IL-17 plays a crucial role in various pathological conditions. It has been demonstrated that IL-17 plays a protective role against opportunist fungal infections [42] and bacterial [43] infections. Patients with impaired IL-17 signaling are more prone to developing chronic mucocutaneous candidiasis [44]. Conversely, dysregulated IL-17 signaling has been linked to the promotion of several autoimmune diseases [45]. Owing to its central role in the inflammatory processes, the IL-17 signaling pathway has become a focal point of clinical research. Numerous molecules that can modulate this pathway are being successfully used in clinical settings. For instance, Secukinumab which is known to block IL-17 signaling [46] is used to treat several autoimmune diseases including plaque psoriasis, psoriatic arthritis and ankylosing spondylitis [47]. As the role of IL-17 signaling in human pathologies, including viral infections and cancer, [48] continues to expand, there is a growing need for novel therapeutics targeting this pathway. Among the emerging technologies, peptide therapy stands out due to its lower toxicity and enhanced specificity. The ongoing preclinical and clinical developments of numerous peptides [49] underscores their potential in therapeutic applications.

In this study, we employed machine learning based approaches to classify IL-17 inducing peptides. Previous research introduced a machine learning prediction tool ‘iL17escan’ which predicted IL-17 inducing peptides with an accuracy of 78.57% and an MCC of 0.57 on the validation dataset [23]. However, a major limitation of that study was the insufficient number of experimentally validated IL-17 inducing peptides. Consequently, we designed this study to integrate the newly validated peptides and improve the prediction accuracy of IL-17 inducing peptides. In comparison to the previous study which included 338 experimentally validated peptides, our research incorporated 618 experimentally validated IL-17 inducing peptides.

The preliminary analysis of the compositional analysis of the peptides revealed significant enrichment of certain amino acids including Leucine, Arginine, Serine, and Tryptophan in the IL-17 inducing peptides (p-value < 0.01). Additionally, dipeptide analysis which represents the pairwise occurrence of amino acids revealed the enrichment of certain amino acids pairs in the IL-17 inducing peptides. For instance dipeptides including “AL”, “LK”, “PL”, “LS”, “IL”, “LR”, “RL”, “IS” and “SV” were significantly (p-value < 0.01) enriched in the IL-17 inducing peptides. These results indicate that the amino acids and dipeptides that are differentially enriched in the positive and the negative datasets might be helpful towards their classification. Furthermore, we used the sequence alignment methods as well to improve upon the prediction of the IL-17 inducing peptides. We employed the widely used alignment-based method “BLAST” to classify IL-17 and non-IL-17 inducers. This approach involves searching the query peptide sequence in a database of known IL-17 and non-IL-17 inducers. If a match is found, the annotation of the database sequence is assigned to the query leading to the accurate prediction of the IL-17 inducing peptides.

Apart from the peptide composition and BLAST, using the MERCI algorithm, we identified motifs such as PVI, FSR, FSRV and NPVI, which were exclusively present in IL-17 inducers, indicating their potential contribution towards the classification of IL-17 inducing peptides. We utilized the Pfeature algorithm to extract features from 16 descriptors of the compositional module for both the positive and negative datasets, generating a total of 9151 features for each peptide sequence. Prediction models were developed for each descriptor using machine learning classifiers, including k-Nearest Neighbour (kNN), Random Forest (RF) and Support Vector Machine (SVM) with all 9151 features. The threshold value for each model was selected to minimize the difference between the True Positive Rate (TPR) and False Positive Rate (FPR) during cross validation. Notably, the model using DPC features combined with the SVM classifier achieved an MCC of 0.6 during internal validation and 0.63 during external validation. The SVM model built with DPC features outperformed the other models.

To create a more robust prediction model that reduces false positives and false negatives, we leveraged a hybrid approach combining similarity search (BLAST), motif search (MERCI) and alignment-free methods (ML models). This approach overcomes the limitations of individual methods and is commonly used in classification problems. We found that the hybrid approach produced more robust results, with the MCC increasing from 0.6 to 0.65 during internal validation and from 0.63 to 0.68 during external validation. All these models were incorporated in a user-friendly webserver to enable scientists predict the IL-17 inducing potential of the query peptide.

Thus, in this study, we have successfully used the integrated hybrid approach to improve the prediction and accuracy of IL-17 inducing peptides as compared to the existing methods. However, the availability of the more experimentally validated IL-17 inducing peptides is highly desirable as this will help in refining machine learning models leading to better prediction potential.

## Data availability Statement

All the data used in the manuscript is available at www.soodlab.com/iil17pred

## Author Contributions

PA and NP performed the experiments. LA designed and developed the webserver. VS conceived the idea and supervised the study. PA, NP and VS wrote the first draft. BK helped in all the experiments, supervised the study and finalized the draft. All authors have read and approved the final manuscript.

## Supplementary information

Supplementary data is available online.

## Acknowledgements

PA and BK are thankful to Hansraj college for the start-up grant. VS is thankful to UGC for the FRP and the start-up grant.

