## Supplementary Figure 1 for "Effective prediction of IL-17 inducing peptides using hybrid approach: iIL17pred"

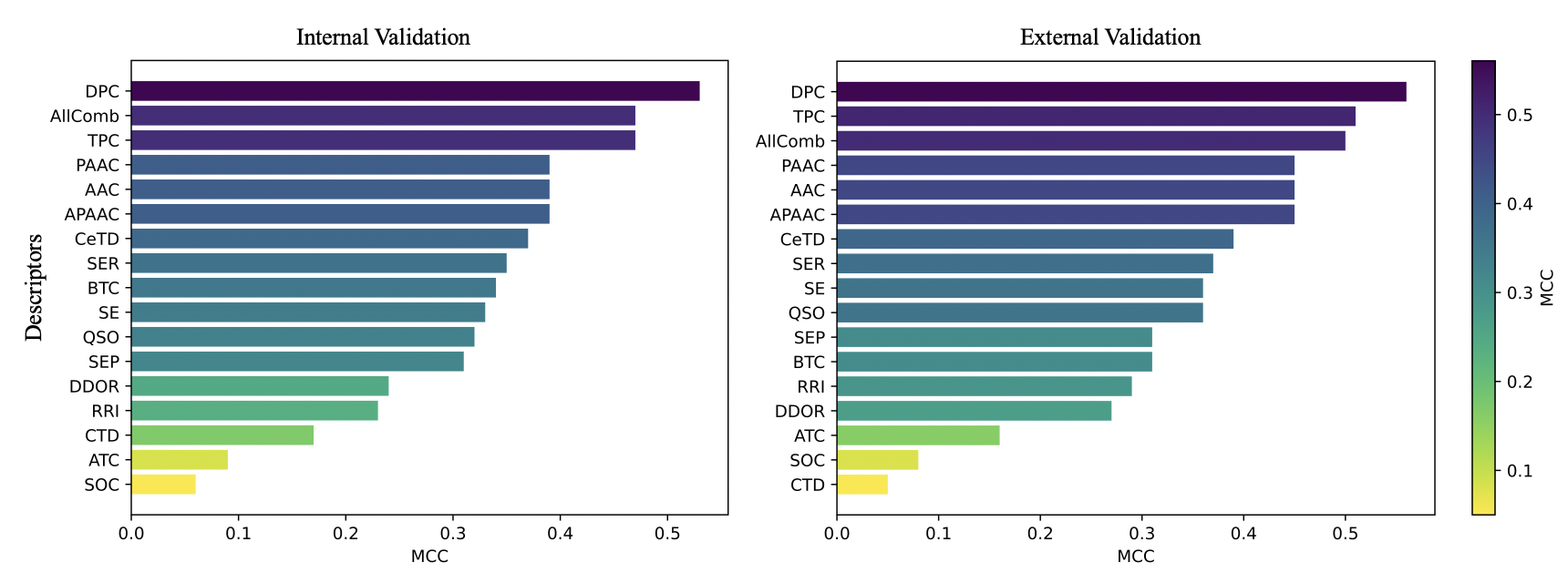

Supplementary Figure 1: The representation of performance metrics of models using different compositional features with k-Nearest Neighbour (kNN) classifier. Matthews Correlation Coefficient (MCC) values are on the x-axis and the features are on the y-axis. The descriptors are ranked based on their MCC values, represented by color gradients. Higher MCC values indicate better performance in predicting outcomes
