## Supplementary Figure 3 for "Effective prediction of IL-17 inducing peptides using hybrid approach: iIL17pred"

Supplementary Figure 3: shows the comparison of the performance of the k-Nearest Neighbor (kNN) based model alone and in combination with BLAST. These models were built on dipeptide composition (DPC) features. The X-axis represents the performance metrics and Y-axis represents the values
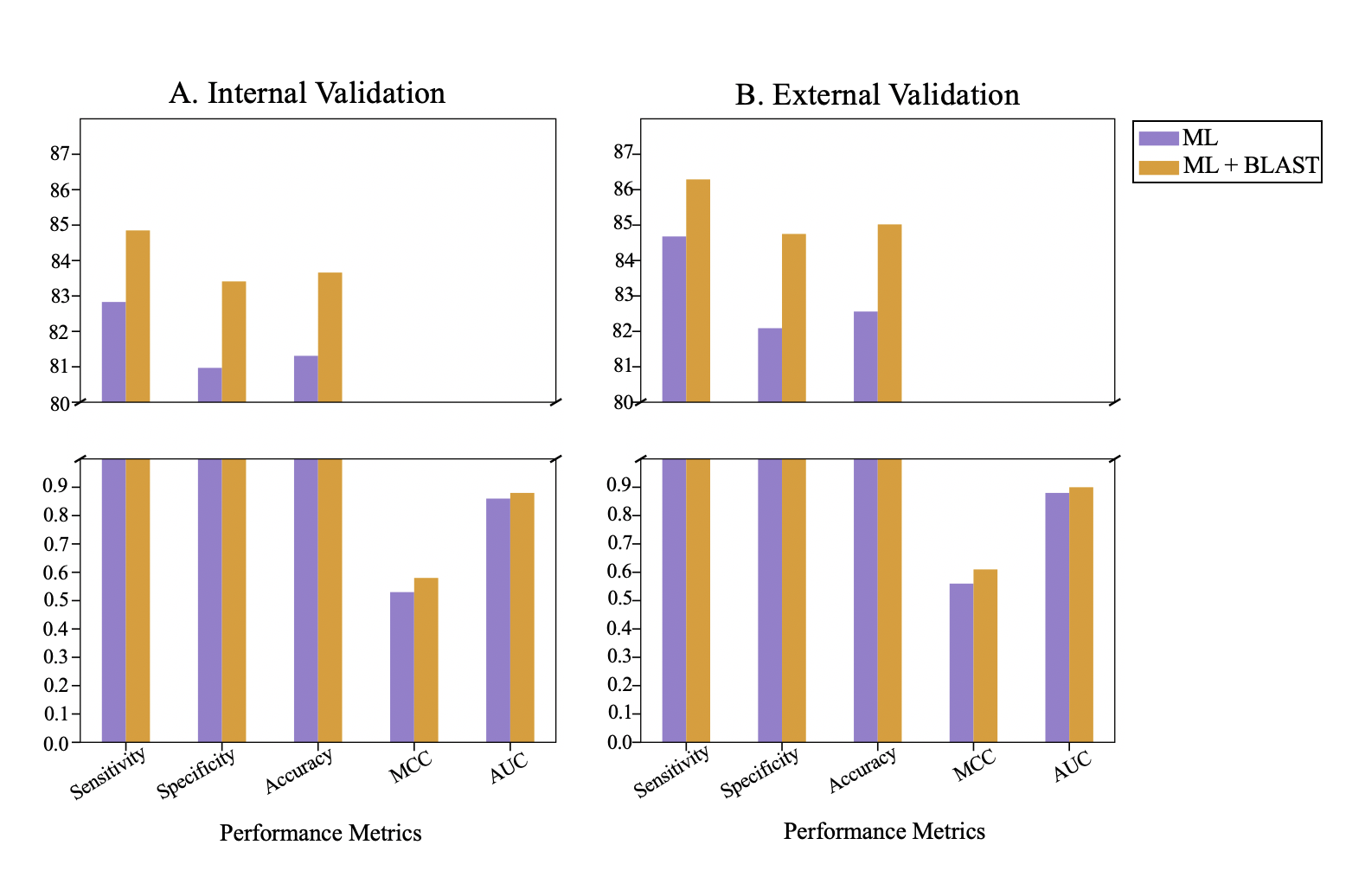
.
