## Supplementary Figure 5 for "Effective prediction of IL-17 inducing peptides using hybrid approach: iIL17pred"

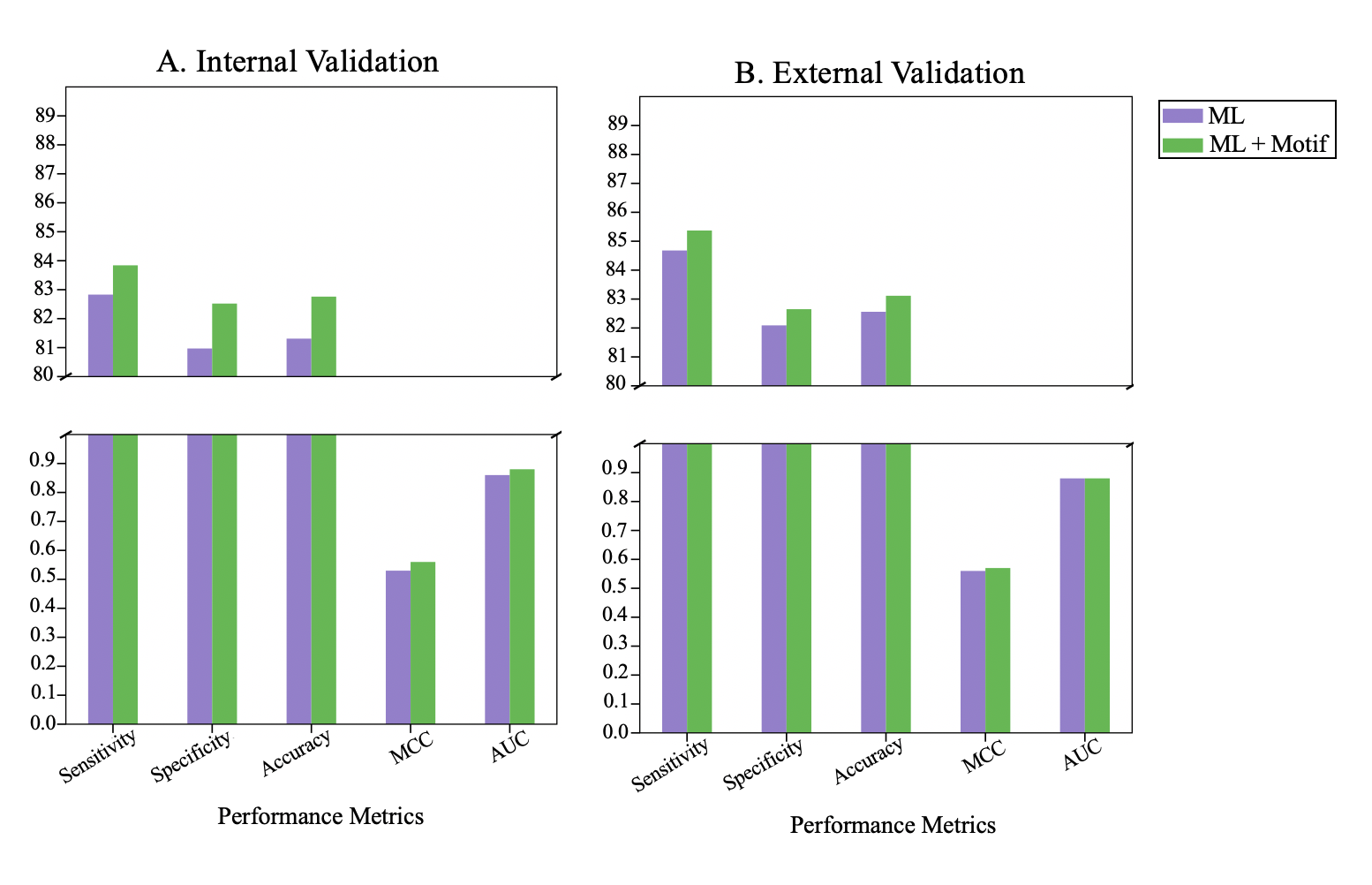

Supplementary Figure 5: shows the comparison of the performance of k-Nearest Neighbor (kNN) based model alone and in combination with MERCI. These models were built on dipeptide composition (DPC) features. The X-axis represents the performance metrics and the Y-axis represents the values.
