## Supplementary Figure 6 for "Effective prediction of IL-17 inducing peptides using hybrid approach: iIL17pred"

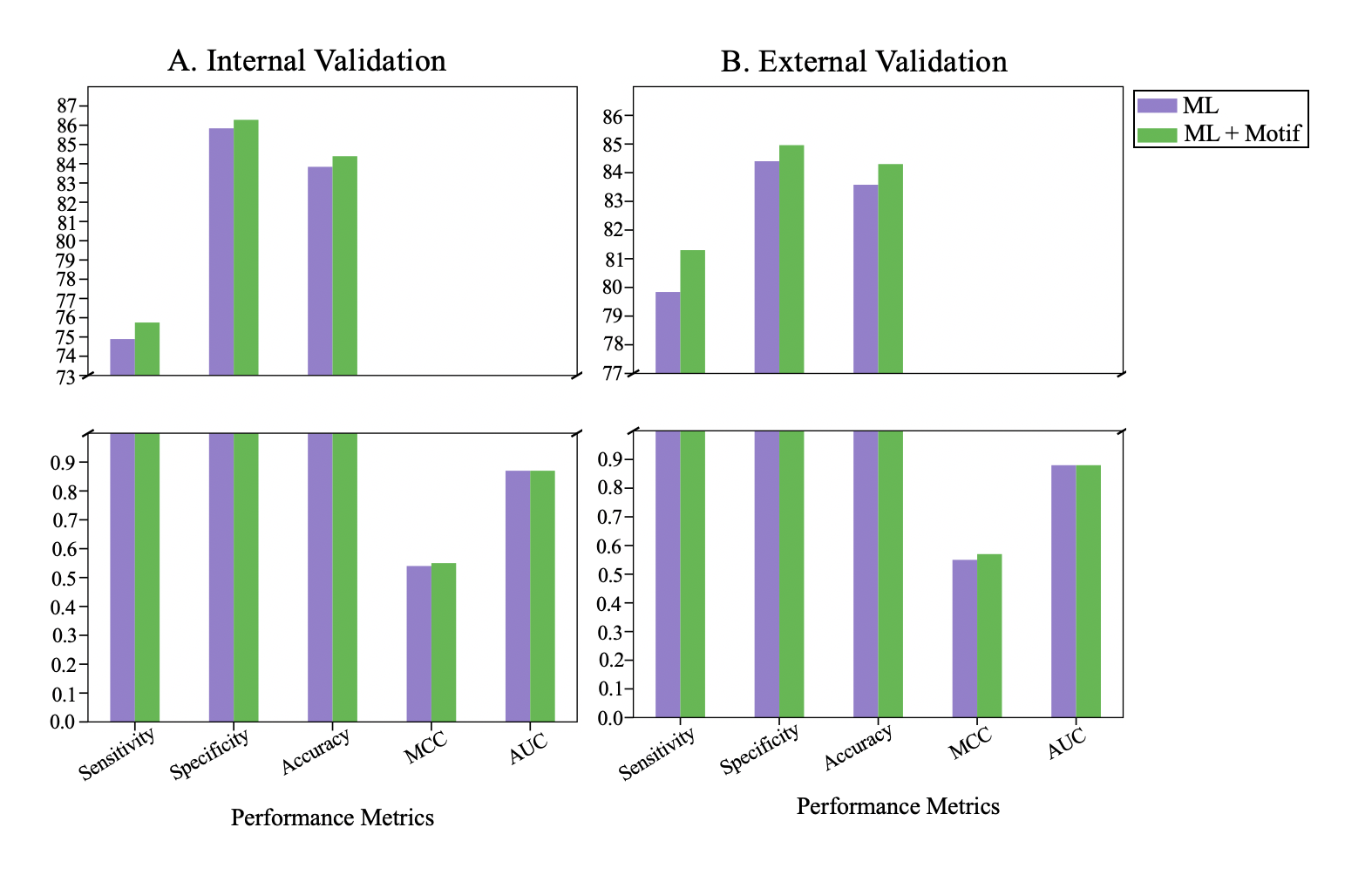

Supplementary Figure 6 shows the comparison of the performance of the Random Forest (RF) based model alone and in combination with MERCI. These models were built on dipeptide composition (DPC) features. The X-axis represents the performance metrics and Y-axis represents the values.
