## Supplementary Figure 7 for "Effective prediction of IL-17 inducing peptides using hybrid approach: iIL17pred"

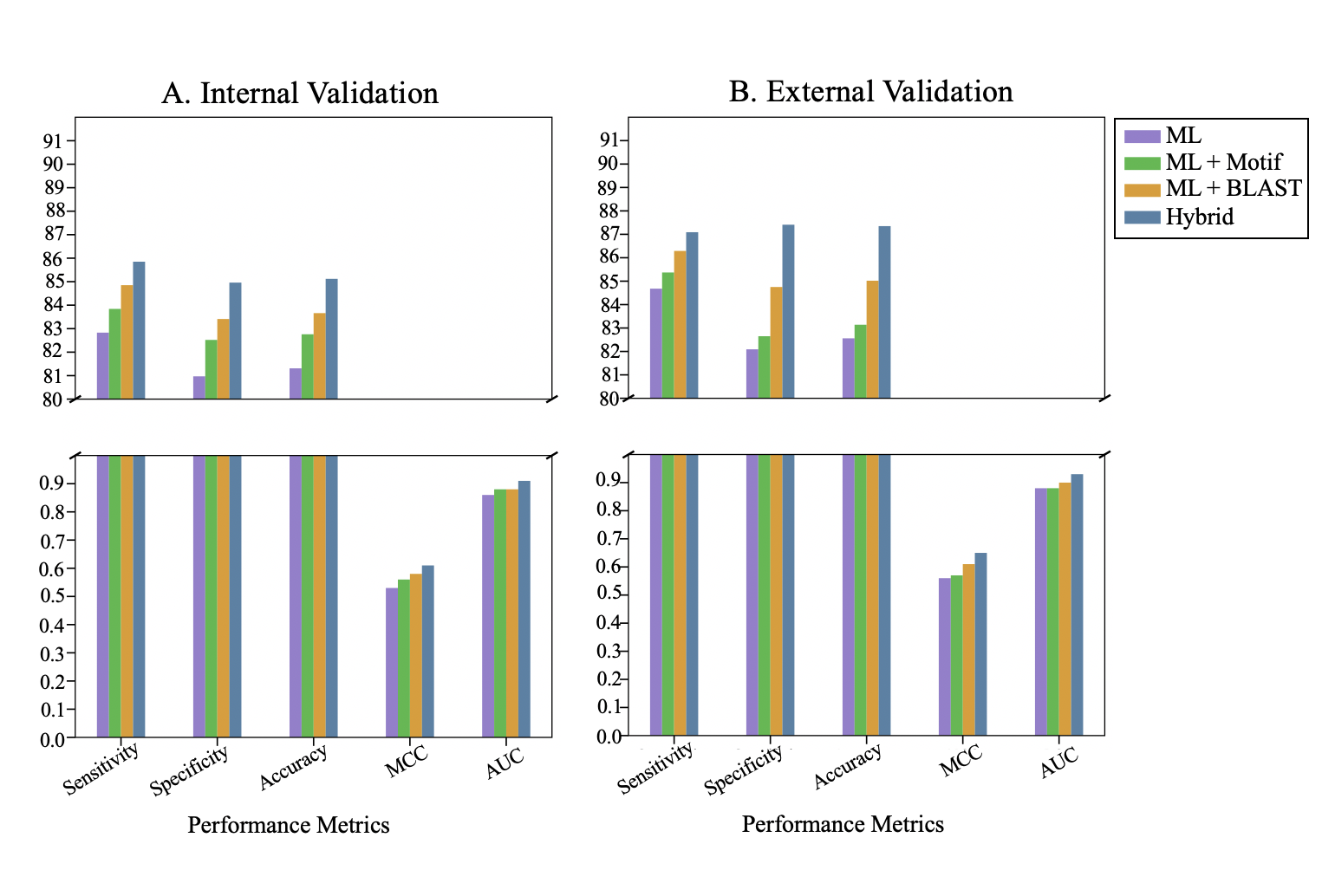

Supplementary Figure 7 shows the comparison of the performance of the k-Nearest Neighbour (kNN) based model in combination with MERCI and BLAST, and the hybrid model. These models were built on DPC features. The X-axis represents the performance metrics and Y-axis represents the values.
