## Supplementary Figure 8 for "Effective prediction of IL-17 inducing peptides using hybrid approach: iIL17pred"

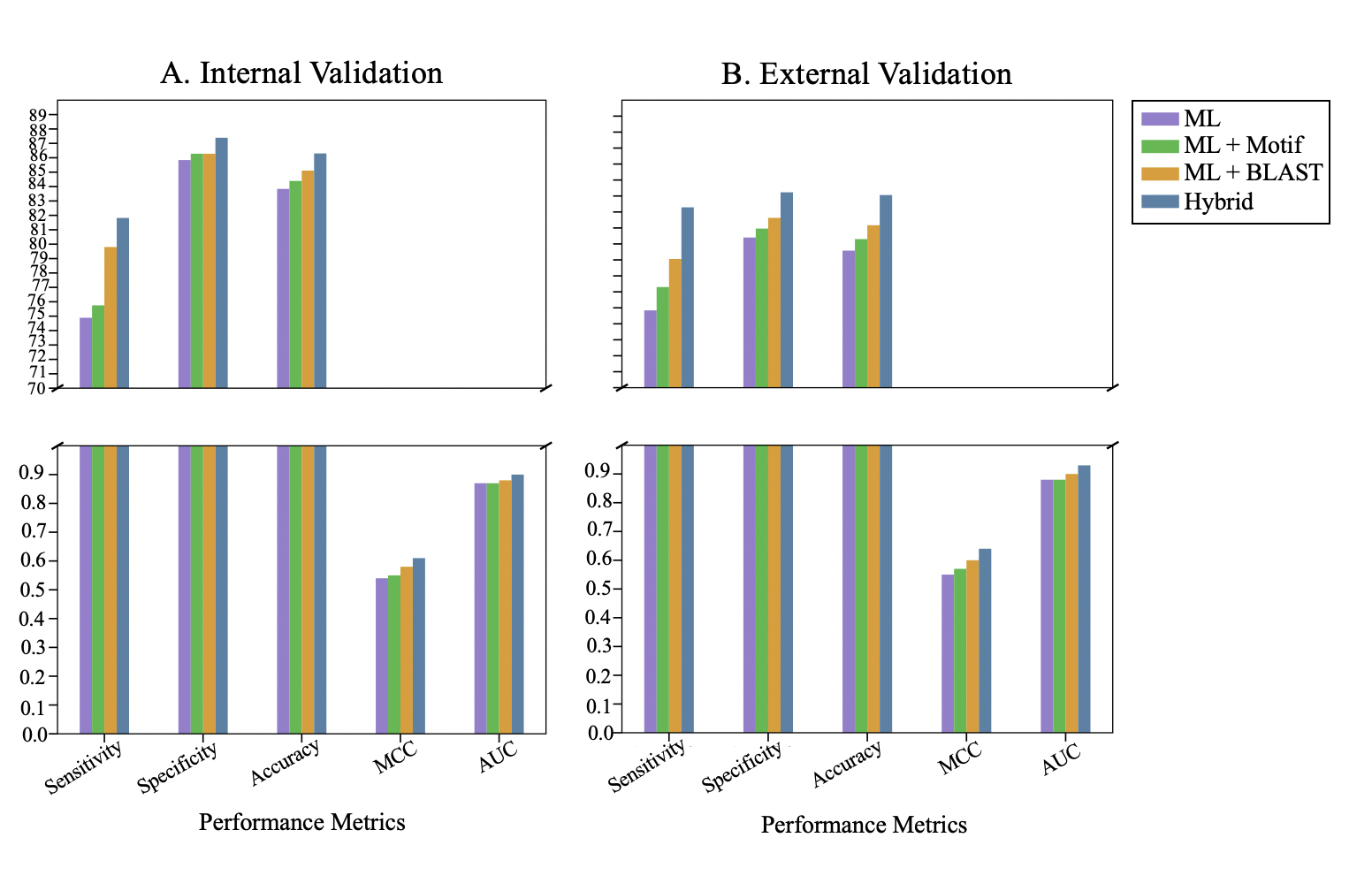

Supplementary Figure 8 shows the comparison of the performance of the Random Forest (RF) based model in combination with MERCI and BLAST, and the hybrid model. These models were built on DPC features. The X-axis represents the performance metrics and the Y-axis represents the values.
