## Supplementary Table 1 for "Effective prediction of IL-17 inducing peptides using hybrid approach: iIL17pred"

**Supplementary Table 1:** A list of descriptors with brief description and the corresponding number of features computed using Pfeature algorithm

| Name of descriptor | Description of descriptor | Number of features |
| --- | --- | --- |
| SE | Shannon-entropy of protein | 1 |
| BTC | Bond composition | 4 |
| ATC | Atomic composition | 5 |
| SOCN | Sequence order coupling number | 6 |
| AAC | Amino acid composition | 20 |
| DDOR | Distance distribution of residue | 20 |
| RRI | Residue repeat information | 20 |
| SER | Shannon entropy of all amino acids | 20 |
| PAAC | Pseudo amino acid composition | 23 |
| SEP | Shannon-entropy of physiochemical property | 25 |
| APAAC | Amphiphilic pseudo amino acid composition | 29 |
| QSO | Quasi-sequence order | 46 |
| CeTD | Composition-enhanced transition distribution | 189 |
| CTD | Conjoint triad calculator of the descriptor | 343 |
| DPC | Dipeptide composition | 400 |
| TPC | Tripeptide composition | 8000 |

Abbreviation: AAC, Amino Acid Composition; DPC, Dipeptide Composition; TPC, Tripeptide Composition; ATC, Atomic Composition; BTC, Bond Composition; DDOR, Distance distribution of residue; RRI, Residue repeat Information; SE, Shannon-entropy of protein; SER, Shannon entropy of all amino acids; SEP, Shannon-entropy of physiochemical property; CTD, Conjoint triad calculation of the descriptor; CeTD, Composition-enhanced transition distribution; PAAC, Pseudo amino acid composition; APAAC, Amphiphilic pseudo amino acid composition; QSO, Quasi-sequence order; SOCN, Sequence order coupling number.
