## Supplementary Table 2 for "Effective prediction of IL-17 inducing peptides using hybrid approach: iIL17pred"

### Supplementary Table 2: Welch’s *t*-test on amino acid frequency among the IL-17 and non-IL-17 inducing peptides.

| **AminoAcid** | **Frequency in IL-17 inducng peptides** | **Frequency in non-IL-17 inducing peptides** | **p value** |
| --- | --- | --- | --- |
| A | 8.3 | 8.85 | 0.14403446 |
| C | 0.97 | 1.19 | 0.08453341 |
| D | 4.19 | 5.77 | 3.06E-10 |
| E | 5.06 | 6.19 | 4.26E-05 |
| F | 4.88 | 4.46 | 0.09333913 |
| G | 6.35 | 7.19 | 0.00671864 |
| H | 2.1 | 2.25 | 0.40985631 |
| I | 5.64 | 5.47 | 0.52641563 |
| K | 5.73 | 6.58 | 0.00219333 |
| L | 11.1 | 8.33 | 1.95E-12 |
| M | 1.76 | 2.03 | 0.08044057 |
| N | 4.8 | 4.49 | 0.2332555 |
| P | 5.15 | 5.42 | 0.34898879 |
| Q | 3.94 | 3.66 | 0.24651232 |
| R | 5.19 | 4.04 | 7.15E-06 |
| S | 7.52 | 6.21 | 8.89E-05 |
| T | 5.25 | 5.57 | 0.23301441 |
| V | 6.36 | 7.22 | 0.00178337 |
| W | 1.46 | 1.09 | 0.00686917 |
| Y | 4.26 | 4.01 | 0.28755691 |
