## Supplementary Table 3 for "Effective prediction of IL-17 inducing peptides using hybrid approach: iIL17pred"

### Supplementary Table 3: Welch’s *t*-test on Dipeptide frequency among the IL-17 inducing and non-IL-17 inducing peptides

| Dipeptide | Frequency in IL-17 inducng peptides | Frequency in non-IL-17 inducing peptides | pvalue |
| --- | --- | --- | --- |
| HD | 0.01 | 0.15 | 5.50E-10 |
| FC | 0 | 0.09 | 7.84E-10 |
| SL | 0.98 | 0.39 | 2.75E-08 |
| GP | 0.27 | 0.64 | 2.14E-06 |
| LL | 1.26 | 0.69 | 2.15E-05 |
| GD | 0.19 | 0.43 | 2.23E-05 |
| HI | 0.06 | 0.2 | 2.29E-05 |
| EF | 0.18 | 0.41 | 3.65E-05 |
| TG | 0.25 | 0.49 | 7.31E-05 |
| DV | 0.25 | 0.5 | 0.00016041 |
| PL | 0.67 | 0.33 | 0.00021191 |
| CP | 0.02 | 0.09 | 0.00025993 |
| AL | 1.17 | 0.75 | 0.00032137 |
| DK | 0.23 | 0.44 | 0.0003429 |
| VV | 0.35 | 0.61 | 0.00046134 |
| ID | 0.18 | 0.37 | 0.00056297 |
| IS | 0.56 | 0.28 | 0.00056767 |
| AC | 0.02 | 0.08 | 0.00061255 |
| VK | 0.27 | 0.47 | 0.00067957 |
| ED | 0.21 | 0.4 | 0.0006988 |
| VD | 0.33 | 0.57 | 0.00076183 |
| RG | 0.19 | 0.37 | 0.00079933 |
| DN | 0.1 | 0.24 | 0.00088504 |
| LK | 0.96 | 0.6 | 0.00093509 |
| WT | 0.18 | 0.03 | 0.00099169 |
| IL | 0.73 | 0.43 | 0.00109026 |
| LS | 0.82 | 0.5 | 0.0011026 |
| CG | 0.05 | 0.14 | 0.00116746 |
| SM | 0.04 | 0.12 | 0.00145025 |
| MN | 0.04 | 0.11 | 0.0016357 |
| KE | 0.23 | 0.42 | 0.00169974 |
| LR | 0.75 | 0.46 | 0.00174355 |
| SV | 0.61 | 0.34 | 0.00194345 |
| TD | 0.13 | 0.28 | 0.0022796 |
| YK | 0.18 | 0.35 | 0.00231864 |
| WS | 0.02 | 0.07 | 0.00246955 |
| FF | 0.25 | 0.08 | 0.00277569 |
| SF | 0.39 | 0.19 | 0.00283321 |
| AD | 0.26 | 0.44 | 0.00283773 |
| YF | 0.38 | 0.19 | 0.00298915 |
| DA | 0.34 | 0.55 | 0.00324774 |
| DD | 0.18 | 0.34 | 0.00337967 |
| CE | 0.02 | 0.09 | 0.00358684 |
| PS | 0.47 | 0.25 | 0.00405597 |
| DH | 0.05 | 0.14 | 0.00405702 |
| EE | 0.27 | 0.48 | 0.00444974 |
| PE | 0.29 | 0.47 | 0.00463285 |
| CM | 0 | 0.02 | 0.0048284 |
| YM | 0.04 | 0.11 | 0.00484001 |
| RS | 0.41 | 0.21 | 0.00486081 |
| RL | 0.58 | 0.36 | 0.00550866 |
| VA | 0.41 | 0.63 | 0.00582278 |
| RP | 0.34 | 0.16 | 0.00597297 |
| RT | 0.31 | 0.14 | 0.00644458 |
| NQ | 0.06 | 0.15 | 0.00644716 |
| AG | 0.48 | 0.7 | 0.00697073 |
| PH | 0.05 | 0.12 | 0.00720022 |
| YG | 0.32 | 0.16 | 0.00748762 |
| EA | 0.39 | 0.58 | 0.0086584 |
| EH | 0.04 | 0.1 | 0.00899286 |
| NL | 0.61 | 0.38 | 0.01060521 |
| NY | 0.15 | 0.28 | 0.01082601 |
| IP | 0.26 | 0.42 | 0.01098834 |
| LI | 0.6 | 0.37 | 0.01251992 |
| IE | 0.21 | 0.36 | 0.01260389 |
| HQ | 0.04 | 0.1 | 0.01303608 |
| WL | 0.22 | 0.09 | 0.01328881 |
| WY | 0.13 | 0.04 | 0.01412663 |
| KG | 0.37 | 0.58 | 0.01413489 |
| KR | 0.35 | 0.19 | 0.01603214 |
| NP | 0.37 | 0.21 | 0.01693738 |
| GK | 0.36 | 0.53 | 0.01743795 |
| KK | 0.28 | 0.44 | 0.01767207 |
| AK | 0.33 | 0.49 | 0.01808907 |
| PG | 0.42 | 0.62 | 0.01889294 |
| YC | 0.02 | 0.07 | 0.0189072 |
| SR | 0.38 | 0.22 | 0.02101081 |
| YA | 0.45 | 0.28 | 0.02296703 |
| SA | 0.71 | 0.5 | 0.02315584 |
| AH | 0.22 | 0.1 | 0.02318017 |
| QW | 0.11 | 0.03 | 0.02465904 |
| MP | 0.04 | 0.11 | 0.02617354 |
| GM | 0.08 | 0.16 | 0.02642723 |
| MS | 0.07 | 0.15 | 0.02701287 |
| NR | 0.23 | 0.12 | 0.02885517 |
| PA | 0.39 | 0.56 | 0.02895995 |
| TV | 0.37 | 0.53 | 0.03040586 |
| YV | 0.15 | 0.25 | 0.03103247 |
| PT | 0.26 | 0.39 | 0.03105641 |
| LM | 0.24 | 0.13 | 0.03620162 |
| TS | 0.48 | 0.31 | 0.03641342 |
| FS | 0.45 | 0.29 | 0.03671242 |
| GT | 0.25 | 0.38 | 0.03781478 |
| GQ | 0.28 | 0.16 | 0.04015168 |
| GA | 0.53 | 0.7 | 0.04051851 |
| YR | 0.26 | 0.13 | 0.04068894 |
| DL | 0.4 | 0.55 | 0.04174623 |
| IQ | 0.18 | 0.28 | 0.0421023 |
| LG | 0.81 | 0.59 | 0.04301746 |
| ND | 0.14 | 0.24 | 0.04307106 |
| RW | 0.09 | 0.03 | 0.04375745 |
| AA | 0.89 | 1.18 | 0.04644171 |
| EY | 0.13 | 0.21 | 0.0468954 |
| TF | 0.17 | 0.26 | 0.04739983 |
| HP | 0.15 | 0.07 | 0.04925763 |
| QF | 0.11 | 0.18 | 0.05165659 |
| AR | 0.45 | 0.3 | 0.05195684 |
| SC | 0.04 | 0.09 | 0.05275734 |
| ME | 0.22 | 0.12 | 0.05329552 |
| QY | 0.19 | 0.1 | 0.05826443 |
| DM | 0.07 | 0.12 | 0.06004685 |
| EW | 0.14 | 0.06 | 0.06106076 |
| IH | 0.11 | 0.18 | 0.06692444 |
| CR | 0.03 | 0.07 | 0.0670705 |
| KT | 0.25 | 0.35 | 0.06715477 |
| RN | 0.36 | 0.24 | 0.06774033 |
| QG | 0.36 | 0.24 | 0.06921262 |
| KY | 0.27 | 0.39 | 0.06940899 |
| NS | 0.36 | 0.24 | 0.07003711 |
| TQ | 0.21 | 0.12 | 0.0721602 |
| QA | 0.29 | 0.19 | 0.07592051 |
| WK | 0.04 | 0.08 | 0.07655399 |
| GW | 0.15 | 0.08 | 0.07710289 |
| KA | 0.5 | 0.65 | 0.07714036 |
| VE | 0.28 | 0.39 | 0.07846332 |
| RQ | 0.22 | 0.13 | 0.0800478 |
| HV | 0.1 | 0.17 | 0.0822267 |
| MG | 0.05 | 0.11 | 0.08245315 |
| DI | 0.19 | 0.29 | 0.08400579 |
| WH | 0 | 0.01 | 0.08612025 |
| RH | 0.09 | 0.15 | 0.08793235 |
| GY | 0.3 | 0.2 | 0.09154463 |
| QP | 0.34 | 0.22 | 0.0944869 |
| FI | 0.27 | 0.17 | 0.09624948 |
| EL | 0.38 | 0.5 | 0.09926881 |
| EC | 0.03 | 0.06 | 0.1049091 |
| FD | 0.22 | 0.31 | 0.10567499 |
| SI | 0.44 | 0.32 | 0.11205751 |
| GF | 0.31 | 0.22 | 0.11233277 |
| CT | 0.04 | 0.08 | 0.11412262 |
| TE | 0.21 | 0.3 | 0.11458863 |
| HE | 0.04 | 0.09 | 0.12852817 |
| VQ | 0.16 | 0.23 | 0.12885683 |
| LH | 0.23 | 0.16 | 0.12919537 |
| DE | 0.28 | 0.37 | 0.13199879 |
| IY | 0.27 | 0.18 | 0.133311 |
| ES | 0.23 | 0.32 | 0.13396129 |
| NM | 0.08 | 0.13 | 0.13553408 |
| SS | 0.58 | 0.76 | 0.13604115 |
| RI | 0.36 | 0.26 | 0.13720502 |
| II | 0.28 | 0.38 | 0.13730091 |
| AT | 0.51 | 0.64 | 0.13856384 |
| WQ | 0.04 | 0.07 | 0.13890448 |
| VY | 0.29 | 0.2 | 0.13937634 |
| SQ | 0.35 | 0.25 | 0.1424891 |
| HW | 0.06 | 0.02 | 0.14468102 |
| LT | 0.58 | 0.45 | 0.14512271 |
| KV | 0.32 | 0.41 | 0.14570827 |
| KQ | 0.16 | 0.23 | 0.14878493 |
| WD | 0.05 | 0.09 | 0.1493179 |
| VG | 0.39 | 0.49 | 0.15056691 |
| YL | 0.37 | 0.27 | 0.16147847 |
| FP | 0.28 | 0.19 | 0.16397403 |
| SW | 0.06 | 0.1 | 0.16612278 |
| AM | 0.18 | 0.25 | 0.16638725 |
| TK | 0.22 | 0.3 | 0.16696847 |
| DS | 0.37 | 0.28 | 0.16716363 |
| QS | 0.2 | 0.27 | 0.16727324 |
| FH | 0.06 | 0.1 | 0.17040352 |
| TW | 0.16 | 0.1 | 0.17080666 |
| CC | 0.06 | 0.02 | 0.17232417 |
| NK | 0.29 | 0.21 | 0.18003049 |
| QR | 0.2 | 0.13 | 0.18065005 |
| QL | 0.3 | 0.39 | 0.18364398 |
| WN | 0.08 | 0.04 | 0.1891882 |
| CY | 0.07 | 0.03 | 0.18968541 |
| GL | 0.65 | 0.53 | 0.1923873 |
| VM | 0.11 | 0.06 | 0.20136483 |
| RY | 0.2 | 0.13 | 0.20320717 |
| ER | 0.31 | 0.24 | 0.20464075 |
| DT | 0.2 | 0.26 | 0.20486893 |
| AY | 0.37 | 0.29 | 0.20715353 |
| AP | 0.32 | 0.41 | 0.2113213 |
| GH | 0.13 | 0.19 | 0.21261863 |
| DG | 0.33 | 0.41 | 0.21345281 |
| LP | 0.5 | 0.4 | 0.21489858 |
| WF | 0.08 | 0.04 | 0.21507418 |
| DY | 0.17 | 0.24 | 0.21868506 |
| QC | 0.01 | 0.03 | 0.21903524 |
| IG | 0.33 | 0.25 | 0.21967714 |
| MR | 0.13 | 0.08 | 0.22184364 |
| FG | 0.28 | 0.21 | 0.22520289 |
| IR | 0.34 | 0.25 | 0.22615775 |
| EM | 0.11 | 0.15 | 0.22712417 |
| SD | 0.3 | 0.23 | 0.23125239 |
| GE | 0.39 | 0.49 | 0.23244994 |
| VR | 0.24 | 0.32 | 0.23544746 |
| MW | 0.04 | 0.01 | 0.23545452 |
| QM | 0.04 | 0.06 | 0.23593302 |
| RC | 0.07 | 0.04 | 0.23771569 |
| FV | 0.27 | 0.34 | 0.24033335 |
| GS | 0.61 | 0.51 | 0.24328304 |
| WA | 0.12 | 0.08 | 0.2467333 |
| HG | 0.2 | 0.14 | 0.24681482 |
| PV | 0.32 | 0.41 | 0.24687117 |
| WG | 0.05 | 0.08 | 0.24696179 |
| VL | 0.68 | 0.58 | 0.25783179 |
| EP | 0.21 | 0.28 | 0.2598347 |
| PK | 0.22 | 0.28 | 0.26100354 |
| YT | 0.19 | 0.25 | 0.26250447 |
| RA | 0.35 | 0.27 | 0.27443189 |
| LA | 0.84 | 0.72 | 0.27585115 |
| IT | 0.27 | 0.34 | 0.2792771 |
| CH | 0.01 | 0.03 | 0.28513737 |
| GN | 0.34 | 0.27 | 0.28606028 |
| CS | 0.09 | 0.05 | 0.28715939 |
| CN | 0.05 | 0.07 | 0.29148642 |
| HL | 0.27 | 0.21 | 0.29231175 |
| YI | 0.26 | 0.2 | 0.29406543 |
| LE | 0.58 | 0.5 | 0.29625273 |
| FN | 0.36 | 0.28 | 0.30311913 |
| PN | 0.27 | 0.21 | 0.30363106 |
| WR | 0.07 | 0.04 | 0.31082978 |
| RV | 0.42 | 0.35 | 0.3149221 |
| PP | 0.27 | 0.36 | 0.3153024 |
| TH | 0.14 | 0.1 | 0.31811369 |
| YD | 0.23 | 0.29 | 0.31862789 |
| KP | 0.25 | 0.31 | 0.32768357 |
| NA | 0.44 | 0.36 | 0.33057456 |
| MQ | 0.11 | 0.08 | 0.33106467 |
| CI | 0.06 | 0.03 | 0.33565323 |
| YS | 0.25 | 0.2 | 0.33612023 |
| MV | 0.11 | 0.15 | 0.33967973 |
| AI | 0.45 | 0.53 | 0.34244926 |
| KW | 0.12 | 0.08 | 0.34549417 |
| MI | 0.1 | 0.14 | 0.3573049 |
| HF | 0.2 | 0.16 | 0.35773729 |
| FA | 0.31 | 0.25 | 0.35986352 |
| GC | 0.06 | 0.09 | 0.36028711 |
| MY | 0.05 | 0.08 | 0.36188759 |
| HC | 0.06 | 0.04 | 0.36461279 |
| PR | 0.26 | 0.21 | 0.36469606 |
| LN | 0.55 | 0.47 | 0.37138531 |
| CL | 0.08 | 0.11 | 0.37258834 |
| KC | 0.04 | 0.06 | 0.37294909 |
| VH | 0.25 | 0.2 | 0.37593608 |
| YN | 0.2 | 0.25 | 0.38374916 |
| RE | 0.17 | 0.22 | 0.38476666 |
| EI | 0.29 | 0.34 | 0.38622199 |
| YH | 0.06 | 0.08 | 0.38739571 |
| LV | 0.61 | 0.54 | 0.39056579 |
| NN | 0.27 | 0.22 | 0.3934181 |
| YE | 0.22 | 0.27 | 0.39525092 |
| NV | 0.28 | 0.34 | 0.40547201 |
| AQ | 0.28 | 0.23 | 0.40961126 |
| LQ | 0.49 | 0.43 | 0.41106907 |
| PW | 0.05 | 0.07 | 0.4118456 |
| FT | 0.32 | 0.27 | 0.41337025 |
| NI | 0.32 | 0.27 | 0.41414868 |
| QH | 0.13 | 0.1 | 0.41503129 |
| EQ | 0.35 | 0.29 | 0.41768984 |
| MT | 0.1 | 0.13 | 0.42300395 |
| VW | 0.09 | 0.12 | 0.42309129 |
| FE | 0.26 | 0.31 | 0.42433475 |
| CK | 0.05 | 0.07 | 0.42495698 |
| KN | 0.32 | 0.27 | 0.42579419 |
| HY | 0.06 | 0.08 | 0.43532102 |
| NH | 0.14 | 0.1 | 0.44429254 |
| HM | 0.02 | 0.03 | 0.45027332 |
| LF | 0.43 | 0.37 | 0.45379788 |
| GI | 0.38 | 0.43 | 0.46043354 |
| VT | 0.41 | 0.47 | 0.46043531 |
| WM | 0.02 | 0.01 | 0.46055892 |
| HS | 0.15 | 0.12 | 0.46097959 |
| GG | 0.38 | 0.43 | 0.46273667 |
| KF | 0.22 | 0.26 | 0.46315903 |
| EV | 0.41 | 0.46 | 0.46463943 |
| VN | 0.22 | 0.27 | 0.46546029 |
| FQ | 0.17 | 0.2 | 0.46951025 |
| KM | 0.14 | 0.11 | 0.471925 |
| WE | 0.13 | 0.1 | 0.4761948 |
| RK | 0.3 | 0.26 | 0.47681644 |
| RD | 0.21 | 0.18 | 0.47999398 |
| SG | 0.45 | 0.51 | 0.48088102 |
| TR | 0.23 | 0.2 | 0.48436853 |
| EK | 0.39 | 0.44 | 0.48786339 |
| AN | 0.4 | 0.35 | 0.48830159 |
| PF | 0.27 | 0.22 | 0.48880878 |
| IC | 0.08 | 0.06 | 0.49168574 |
| LY | 0.41 | 0.36 | 0.49286826 |
| PC | 0.05 | 0.07 | 0.50068969 |
| GR | 0.24 | 0.28 | 0.50583323 |
| SN | 0.31 | 0.27 | 0.50937133 |
| ET | 0.27 | 0.23 | 0.51183269 |
| ST | 0.29 | 0.33 | 0.51212836 |
| QV | 0.23 | 0.26 | 0.51751135 |
| WC | 0.01 | 0.02 | 0.5264166 |
| TI | 0.27 | 0.31 | 0.53090144 |
| IN | 0.33 | 0.29 | 0.53282889 |
| PI | 0.24 | 0.2 | 0.53683547 |
| AW | 0.09 | 0.07 | 0.54546606 |
| IM | 0.13 | 0.11 | 0.54944982 |
| SE | 0.42 | 0.38 | 0.55318878 |
| TT | 0.32 | 0.36 | 0.56221965 |
| QI | 0.24 | 0.27 | 0.57113566 |
| MD | 0.09 | 0.11 | 0.57389674 |
| ML | 0.14 | 0.16 | 0.57841225 |
| IF | 0.19 | 0.17 | 0.58176345 |
| IV | 0.28 | 0.31 | 0.58433701 |
| NT | 0.2 | 0.23 | 0.58745317 |
| TN | 0.2 | 0.17 | 0.5920995 |
| HK | 0.1 | 0.08 | 0.59971013 |
| DF | 0.28 | 0.31 | 0.60601511 |
| AS | 0.51 | 0.55 | 0.60723117 |
| TL | 0.52 | 0.48 | 0.60756744 |
| YQ | 0.17 | 0.15 | 0.61499862 |
| MF | 0.11 | 0.09 | 0.62802705 |
| HR | 0.12 | 0.13 | 0.62935927 |
| MM | 0.04 | 0.05 | 0.63646285 |
| QE | 0.23 | 0.2 | 0.64064045 |
| QT | 0.19 | 0.17 | 0.64313607 |
| FR | 0.2 | 0.22 | 0.64813254 |
| FK | 0.37 | 0.34 | 0.66041238 |
| NG | 0.36 | 0.39 | 0.66142974 |
| EG | 0.32 | 0.34 | 0.66161252 |
| SH | 0.13 | 0.15 | 0.66486154 |
| RM | 0.1 | 0.08 | 0.67011564 |
| CD | 0.05 | 0.05 | 0.67842584 |
| MH | 0.04 | 0.03 | 0.68030763 |
| PM | 0.08 | 0.06 | 0.68208461 |
| MC | 0.01 | 0.01 | 0.68547329 |
| TP | 0.42 | 0.38 | 0.68990401 |
| KL | 0.72 | 0.68 | 0.69486746 |
| IA | 0.36 | 0.38 | 0.69974894 |
| VS | 0.52 | 0.49 | 0.71184342 |
| RF | 0.21 | 0.23 | 0.71216618 |
| HA | 0.19 | 0.17 | 0.7161523 |
| VP | 0.33 | 0.35 | 0.71961385 |
| CF | 0.05 | 0.04 | 0.72568156 |
| AV | 0.65 | 0.68 | 0.73173385 |
| RR | 0.21 | 0.23 | 0.73375508 |
| YW | 0.05 | 0.04 | 0.73673364 |
| YY | 0.28 | 0.26 | 0.73709565 |
| TM | 0.1 | 0.09 | 0.74034225 |
| NF | 0.28 | 0.31 | 0.74228831 |
| SK | 0.39 | 0.41 | 0.74775058 |
| QQ | 0.2 | 0.18 | 0.74949751 |
| DW | 0.04 | 0.04 | 0.75300779 |
| IW | 0.04 | 0.04 | 0.75409856 |
| WP | 0.05 | 0.04 | 0.7574878 |
| HT | 0.12 | 0.1 | 0.76359473 |
| WW | 0.02 | 0.02 | 0.77104929 |
| CV | 0.08 | 0.07 | 0.77441072 |
| SY | 0.31 | 0.29 | 0.78295024 |
| PD | 0.19 | 0.17 | 0.78374959 |
| SP | 0.37 | 0.35 | 0.78602752 |
| LD | 0.46 | 0.48 | 0.80629182 |
| TA | 0.53 | 0.51 | 0.80670614 |
| QN | 0.14 | 0.15 | 0.80894327 |
| CA | 0.07 | 0.06 | 0.81377124 |
| GV | 0.46 | 0.44 | 0.81640719 |
| HH | 0.11 | 0.09 | 0.8179808 |
| VF | 0.31 | 0.29 | 0.82529266 |
| KI | 0.34 | 0.33 | 0.83076536 |
| DQ | 0.16 | 0.15 | 0.83809291 |
| MK | 0.11 | 0.12 | 0.84069691 |
| FY | 0.2 | 0.21 | 0.84241498 |
| NC | 0.07 | 0.06 | 0.84620667 |
| HN | 0.09 | 0.09 | 0.85998227 |
| QK | 0.26 | 0.27 | 0.8612943 |
| WI | 0.08 | 0.07 | 0.86200385 |
| VI | 0.38 | 0.39 | 0.86215482 |
| IK | 0.34 | 0.35 | 0.86327833 |
| VC | 0.08 | 0.08 | 0.86937538 |
| AF | 0.41 | 0.42 | 0.87029074 |
| DP | 0.27 | 0.26 | 0.87100722 |
| DC | 0.05 | 0.06 | 0.8730738 |
| EN | 0.28 | 0.27 | 0.8780116 |
| NE | 0.22 | 0.23 | 0.87859115 |
| TY | 0.25 | 0.26 | 0.88199828 |
| FM | 0.09 | 0.09 | 0.89133625 |
| QD | 0.2 | 0.21 | 0.90587741 |
| MA | 0.19 | 0.19 | 0.90587763 |
| CW | 0.02 | 0.02 | 0.90625357 |
| FW | 0.04 | 0.04 | 0.90900833 |
| YP | 0.18 | 0.19 | 0.90927927 |
| LW | 0.05 | 0.05 | 0.91050471 |
| KH | 0.1 | 0.11 | 0.91236435 |
| DR | 0.21 | 0.21 | 0.91313777 |
| KD | 0.39 | 0.39 | 0.91894702 |
| AE | 0.42 | 0.42 | 0.92495707 |
| FL | 0.5 | 0.5 | 0.92953473 |
| KS | 0.34 | 0.34 | 0.93918965 |
| WV | 0.07 | 0.07 | 0.94780254 |
| CQ | 0.03 | 0.03 | 0.95548077 |
| PQ | 0.18 | 0.18 | 0.96594089 |
| TC | 0.1 | 0.09 | 0.97915247 |
| LC | 0.09 | 0.09 | 0.98274287 |
| PY | 0.19 | 0.19 | 0.99395793 |
| NW | 0.06 | 0.06 | 0.99401257 |
