## Supplementary Table 4 for "Effective prediction of IL-17 inducing peptides using hybrid approach: iIL17pred"

### Supplementary Table 4: The distinct motifs present in positive (IL-17 inducing peptides) and negative training data (non-IL-17 inducing peptides)

| S.No | Motifs (IL-17 inducers) | No. of sequence | Motifs (Non-IL-17 inducers) | No. of sequence |
| --- | --- | --- | --- | --- |
| 1 | P V I | 10 | G P A | 28 |
| 2 | F S R | 8 | F C | 27 |
| 3 | F S R V | 8 | A G P | 25 |
| 4 | N P V I | 7 | D V D | 24 |
| 5 | P V I L | 7 | D A L | 23 |
| 6 | F N P V | 6 | E D L | 23 |
| 7 | F N P V I | 6 | E K G | 23 |
| 8 | F N P V I L | 6 | V D V | 23 |
| 9 | F N P V I L R | 6 | E I D | 22 |
| 10 | F N P V I L R P | 6 | K A A | 22 |
| 11 | I L R P | 6 | A E V | 21 |
| 12 | N P V I L | 6 |  |  |
| 13 | N P V I L R | 6 |  |  |
| 14 | N P V I L R P | 6 |  |  |
| 15 | P V I L R | 6 |  |  |
| 16 | P V I L R P | 6 |  |  |
| 17 | V I L R | 6 |  |  |
| 18 | V I L R P | 6 |  |  |
| 19 | Y F N P | 6 |  |  |
| 20 | Y F N P V | 6 |  |  |
| 21 | Y F N P V I | 6 |  |  |
| 22 | Y F N P V I L | 6 |  |  |
| 23 | Y F N P V I L R | 6 |  |  |
| 24 | Y F N P V I L R P | 6 |  |  |
