## Supplementary Table 5 for "Effective prediction of IL-17 inducing peptides using hybrid approach: iIL17pred"

**Supplementary Table 5**: Performance-metrics of machine learning based models on internal and external validation developed by utilizing various compositional features in conjunction with k-nearest neighbour (kNN) classifier. The maximum values in each performance metrics are highlighted in bold.

| Descriptor | No. of features | Sensitivity | Specificity | Accuracy | MCC | ROC | Sensitivity | Specificity | Accuracy | MCC | ROC |
| --- | --- | --- | --- | --- | --- | --- | --- | --- | --- | --- | --- |
|  |  | Internal validation | | | | | External validation | | | |  |
| AAC | 20 | 73.69 | 74.62 | 74.46 | 0.39 | 0.79 | 79.03 | 76.24 | 76.74 | 0.45 | 0.82 |
| DPC | 400 | **82.83** | **80.97** | **81.31** | **0.53** | **0.86** | **84.68** | **82.09** | **82.56** | **0.56** | **0.88** |
| TPC | 8000 | 75.91 | 80.69 | 79.83 | 0.47 | 0.81 | 77.41 | 82.62 | 81.68 | 0.51 | 0.82 |
| ATC | 5 | 53.04 | 58.19 | 57.26 | 0.09 | 0.58 | 60.48 | 60.64 | 60.61 | 0.16 | 0.62 |
| BTC | 4 | 71.06 | 71.03 | 71.04 | 0.34 | 0.76 | 68.55 | 70.03 | 69.77 | 0.31 | 0.74 |
| DDOR | 20 | 60.93 | 69.13 | 67.66 | 0.24 | 0.68 | 64.52 | 69.68 | 68.75 | 0.27 | 0.73 |
| RRI | 20 | 66.59 | 62.62 | 63.34 | 0.23 | 0.69 | 70.16 | 66.31 | 67.00 | 0.29 | 0.70 |
| SE | 1 | 69.23 | 71.16 | 70.82 | 0.33 | 0.75 | 72.58 | 72.16 | 72.24 | 0.36 | 0.78 |
| SER | 20 | 71.06 | 72.14 | 71.95 | 0.35 | 0.78 | 72.58 | 72.87 | 72.82 | 0.37 | 0.78 |
| SEP | 25 | 72.47 | 67.13 | 69 | 0.31 | 0.75 | 70.16 | 69.14 | 69.33 | 0.31 | 0.72 |
| CTD | 343 | 70.04 | 52.04 | 55.27 | 0.17 | 0.65 | 54.03 | 52.12 | 52.47 | 0.05 | 0.53 |
| CeTD | 189 | 72.87 | 73.43 | 73.33 | 0.37 | 0.79 | 73.39 | 74.29 | 74.13 | 0.39 | 0.79 |
| PAAC | 23 | 72.88 | 74.58 | 74.27 | 0.39 | 0.79 | 79.03 | 76.06 | 76.6 | 0.45 | 0.82 |
| APAAC | 29 | 73.88 | 74.71 | 74.56 | 0.39 | 0.8 | 80.65 | 75.0 | 76.01 | 0.45 | 0.83 |
| QSO | 46 | 67.60 | 71.70 | 70.97 | 0.32 | 0.74 | 70.97 | 73.40 | 72.97 | 0.36 | 0.75 |
| SOC | 6 | 52.42 | 53.81 | 53.56 | 0.06 | 0.52 | 57.26 | 53.01 | 53.78 | 0.08 | 0.53 |
| All_Comb(D) | 9151 | 82.79 | 74.58 | 76.05 | 0.47 | 0.83 | 83.06 | 78.36 | 79.22 | 0.5 | 0.84 |
